# Palmdelphin facilitates R-spondin2 secretion to activate Wnt signaling and promote colorectal cancer stemness and tumorigenesis

**DOI:** 10.1101/2025.04.03.646911

**Authors:** Yuning Yang, Jinsen Shi, Yuping Yang, Sihan Liu, Yi Li, Lian Feng, Rui Yan, Jiannan Yao, Lushan Chen, Ling Ding, Zhuqing Zhang, Hailan Feng, Hong Chen, Qin Lu, Ting Yan, Zixing Yan, Parthasarathy Chandrakesan, Dongfeng Qu, Jian Du, Zhiyun Cao, Jun Peng, Nathaniel Weygant

**Affiliations:** Academy of Integrative Medicine, Fujian Univ. of TCM, Fuzhou, China; Dept. of Oncology, Beijing-Chao Yang Hospital, Beijing, China; Dept. of Pathology, Fuzhou Union Hospital, Fuzhou, China; Dept. of Integrative Medicine, Shanghai Jiaotong University, Shanghai, China; Second Affiliated Hospital of Fujian Univ. of TCM, Fuzhou, China; Affiliated Fuzhou Hospital, Fujian Univ. of TCM, Fuzhou, China; Core Environmental Monitoring Lab, Harrisburg, USA; Dept. of Medicine, University of Oklahoma Health Sciences Center, Oklahoma City, USA

**Author notes:** **Materials and Correspondence:** Nathaniel Weygant PhD, Jun Peng PhD, Zhiyun Cao PhD. Equal First Authors (Yuning Yang, Jinsen Shi).

**Keywords:** Palmdelphin, Patient-derived organoid, Cancer stem cell, R-spondin2, Wnt signaling pathway, Colorectal cancer

## Abstract

The Wnt signaling pathway is a key driver of stemness and progression which contributes to mortality in colorectal cancer (CRC). R-spondins bind to LGR receptors to inhibit ubiquitin E3 ligases, thus protecting Frizzled from degradation and activating downstream Wnt signaling. Herein, we identify Palmdelphin (*PALMD*) as a functional marker of CRC stem cells, interspersed between intestinal and colonic crypt base epithelial cells, and predictive of aggressive CMS4 CRC and poor survival. Gene knockdown and overexpression studies revealed that *PALMD* initiates paracrine activation of Wnt/β-catenin signaling via interacting with and facilitating the secretion of R-spondin2 (*RSPO2*), resulting in enhanced stemness and tumor growth *in vitro* and *in vivo*. Physiologic or pharmacologic inhibition of the *PALMD-RSPO2* axis using R-spondin2-specific antibody or 6-methyl-1,3,8-trihydroxyanthraquinone (emodin), respectively, attenuates *PALMD-* mediated Wnt reporter activation, self-renewal, and tumorigenesis in cell and patient-derived organoid models. Together, these findings identify *PALMD* as a previously unknown player in Wnt signaling in CRC, and underscore the pro-tumorigenic role of R-spondin2 in this context.

## Introduction

Colorectal cancer (CRC) is the second most lethal tumor worldwide^1^. Screening methods for CRC are constantly improving and relevant first-line treatment schemes such as surgery, chemotherapy, and targeted therapy are utilized to great benefit in patients with early disease^2^. However, a majority of CRCs are discovered at an advanced stage and high rates of drug resistance, recurrence, and metastasis are an important cause of mortality^3,4^. Since 2015, a consensus molecular subtyping (CMS) system has emerged as a tool to improve classification and treatment of CRCs based on pathological and molecular characteristics. The CMS system identifies four major subtypes of CRC including hypermutant CRCs with microsatellite instability (CMS1), Wnt/MYC-driven and well-differentiated epithelial-like CRCs (CMS2), metabolically dysregulated epithelial-like CRCs (CMS3), and an epithelial-mesenchymal transition (EMT) and cancer stem cell (CSC)-associated mesenchymal-like CRC subtype (CMS4). Among these four subtypes, CMS4 CRCs are the most aggressive with the worst overall and relapse-free survival, while CMS1 CRCs demonstrate the poorest survival following relapse^5^. Overall, a better understanding of CRC etiology and the development of new therapies that can target aggressive CRC subtypes may lead to improvements in patient survival.

CRCs arise from the intestinal epithelium which is structured to maintain homeostasis in the presence of inflammation through specialized sensory and protective cells and highly responsive stem cells^6,7^. Dynamic components within this system sense changes in the microenvironment and communicate with the immune system, which in turn reprograms intestinal stem cells (ISCs) to formulate a proliferative response^6–9^. Together these components repair the epithelium, but may also be a source of tumor initiation due to stem cells and long-lived tuft cells harboring mutations^10^. ISCs give rise to tumors and together with their progeny can continuously replace the tumor epithelium, which is a key factor in tumor progression^11–13^.

ISCs, the first stem cells to be definitively identified in solid tissue, were originally shown to be regulated by Wnt signaling and present in the intestinal crypt^14^. Using Cre recombinase-based transgenic mouse technology, the Clevers group identified ISCs as Lgr5+ cells interspersed between Paneth cells in the epithelial crypt base with ability repopulate the intestinal epithelium in rapid fashion^15^. Subsequent studies confirmed that common CRC alterations such as APC mutation and β-catenin cleavage/nuclear translocation in these cells would initiate tumors and give rise to CSCs that continuously fuel the growth and progression of the tumor^12,13^. Despite confirmation of the role of ISCs and their CSC progeny in human CRCs and extensive attempts at developing targeted therapies against them, a breakthrough has yet to materialize. Therefore, new targets for ISC-mediated tumorigenesis and CSC-mediated progression remain highly sought after.

Palmdelphin (PALMD) is part of the paralemmin phosphoprotein family, the members of which are abundantly distributed in the brain, kidney and other tissues and closely associated with the plasma membrane of brain synapses^16,17^. Paralemmins have been extensively studied in cardiac, immunologic, and neurobiological models^18–21^, but their role in cancer is relatively unexplored. PALM1 is expressed in breast cancer cell lines and tissue, and shows higher expression in invasive ductal carcinomas and estrogen receptor+ subtypes^22^. PALM2 is upregulated and predicts poor prognosis in esophageal squamous cancer (ESCC). Its prenylated form is localized to the cell membrane, and its expression enhances invasive properties of ESCC via interaction with ezrin^23^. Palmdelphin (PALMD) was identified by chromatin immunoprecipitation (ChIP) sequencing as a significant target of phosphorylated p53 (serine 46) in osteosarcoma, where it binds to PALMD’s promoter region and regulates its expression. Adriamycin-induced DNA damage significantly increases PALMD expression in p53-proficient osteosarcoma cells, which enhances apoptosis in this model^24^. More recently, PALMD was identified as an inhibitor of invasion and migration under the control of the ZNF263 transcription factor in uveal melanoma^25^ and an inhibitor of proliferation via the PI3K/AKT pathway in breast cancer^26^.

The Wnt signaling pathway is a lynchpin for stemness and tumor growth in CRC. In canonical Wnt signaling, R-spondins1-4 bind to LGR4-6, which in complex suppress the activity of ubiquitin ligases ZNRF3/RNF43, in effect preventing the degradation of Frizzled and allowing downstream signaling to induce β-catenin nuclear translocation and TCF/LEF-mediated gene transcription. Among R-spondins, R-spondin2 (RSPO2) has the strongest affinity for gastrointestinal epithelial stem cell marker LGR5^27^. Notably, studies using various methodologies have revealed conflicting findings in terms of R-spondin2’s activity in cancer. For instance, Wu *et al.* showed that it may act as a tumor suppressor when expressed in select CRC cell lines^28^, whereas Chartier *et al.* demonstrated that blockade of R-spondin2 using specific antibody potently inhibits CRC patient-derived tumor growth *in vivo*^29^. Additionally, RSPO2 fusions are reported in a subset of human CRCs, mutually exclusive with *APC* mutation, and sufficient to initiate Wnt-mediated tumorigenesis in mouse models^30–32^. In this study, we highlight the role of R-spondin2 paracrine signaling initiated from PALMD expressing CRC cells. Namely, we establish that PALMD serves as a novel marker of CRC stem cells, which activates Wnt signaling via facilitating R-spondin2 secretion. This process fosters CRC stemness and tumorigenesis both *in vitro* and *in vivo*. Thus, our findings underscore the pro-tumor role of R-spondin2 in the context of *PALMD* expression.

## Results

### PALMD is expressed in intestinal crypt base cells and CRC stem cells and predicts prognosis

In order to identify novel CRC stem cell markers, we pursued a combined bioinformatics and immunohistochemistry strategy (**Fig S1A**) based on the Batlle colon ISC signature^33^, while controlling for the influence of disease stage. Our analysis identified significantly increased expression of *PALMD* in high ISC signature colon cancer (**Fig 1A**), strong expression in crypt base epithelial cells consistent with ISCs (**Fig 1B**), and overexpression in CRC tissue (**Fig 1C**, **Fig S1B**). Statistically significant overexpression of *PALMD* was present in CRC epithelium compared to para-cancerous tissue in stage I-II patients as quantified by an experienced pathologist, while *PALMD* expression in tumor stroma was significantly decreased (**Fig 1C-D**, **Fig S1C**). Interestingly, pathological scoring indicated that *PALMD* expression is mostly lost in CRC endothelial and smooth muscle cells, while it is expressed in cancer-associated fibroblasts. We further assessed *PALMD* expression by IHC in advanced (stage III-IV) CRC surgical specimens and found a range of expression patterns from weak, diffusely positive to strong, specific positive staining, which leads us to conclude that *PALMD* is expressed in all stages of CRC (**Fig S1D**).

**Figure 1.**
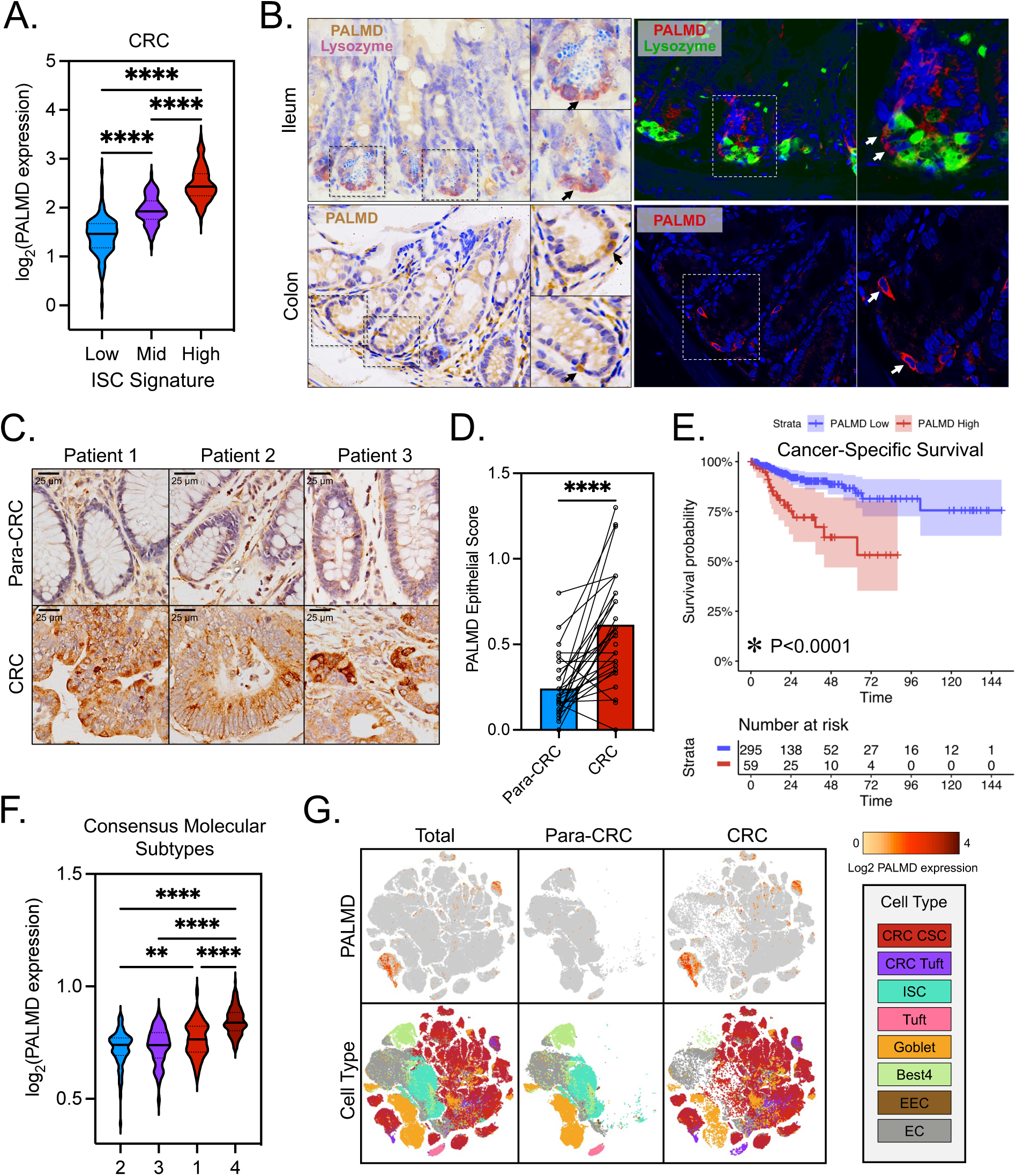
Palmdelphin (*PALMD*) is a cancer stem cell-associated marker in colorectal cancer. **A.** *PALMD* is expressed at progressively higher levels in colon adenocarcinoma according to intestinal stem cell (ISC) signature category (P<0.0001 for all comparisons). **B.** Immunohistochemical and immunofluorescence staining of *PALMD* and Lyzoszyme (Paneth cell marker) in normal C57/B6 moues intestine and colon showing specific expression of *PALMD* in the crypt base of the intestinal and colonic epithelium. **C.** Immunohistochemical staining of *PALMD* in a tissue microarray demonstrating significant overexpression in CRC epithelia compared to adjacent normal (para-CRC) epithelia. **D.** Comparison of pathologist epithelial scoring (intensity x area) for *PALMD* in CRC compared to para-CRC showing a significant increase in CRC (paired T-test P<0.0001). **E.** High expression levels of *PALMD* are associated with significantly reduced cancer-specific survival (CSS) in the TCGA colorectal cancer dataset (log-rank test P<0.0001). **F.** *PALMD* is most strongly expressed in the consensus molecular subtype (CMS) 4 followed by CMS1 (Mann-Whitney U Test: ****P<0.0001, **P<0.01). **G.** t-SNE projection from single-cell RNA sequencing data obtained from the human colon cancer atlas (HCCA) confirming *PALMD* overexpression in CRC compared to para-CRC epithelium, and demonstrating its localization to primarily ISCs and tuft cells in normal epithelium, and CSCs and CRC tuft cells in conditions of cancer.

To further understand *PALMD* in terms of CRC pathology, we sought to assess its association with CRC patient survival, response to therapy, and CRC molecular subtype. Analysis of cancer-specific survival (CSS) and progression-free survival (PFS) in CRC patients from TCGA’s COADREAD dataset revealed that high expression of *PALMD* predicts reduced CSS and PFS (P<0.0001)(**Fig 1E**, **Fig S1E**). To assess this trend in the context of drug resistance, which is associated with CSCs, we selected CRC patients who received first-line chemotherapy and analyzed survival. Among CRC patients receiving first-line chemotherapy, those expressing high levels of *PALMD* had significantly reduced PFS compared to their counterparts (P=0.01)(**Fig S1F**). Molecular profiling analysis for CRC subtypes revealed that *PALMD* was most highly expressed in aggressive EMT/CSC-linked CMS4 tumors, followed by the microsatellite instable CMS1 tumors in the TCGA COAD dataset (**Fig 1F**).

To gain further insight into *PALMD* in the cellular context of CRC, we turned to single-cell RNA sequencing (scRNA-Seq) analysis. Using epithelial data from the Human Colon Cancer Atlas (HCCA), we performed correlation analysis with known cell markers and visualized scRNA-Seq results as a t-SNE projection to further understand *PALMD*+ cell localization in the normal colon and CRC epithelium. Correlation analysis with known stem cell and tuft cell markers in CRC epithelium revealed *PALMD* coexpression with *LGR5*, *NANOG*, and *MYC* in tumor stem/TA-like epithelium, and tuft cell speciation factor *POU2F3* and pluripotency factors *SOX2*, *NANOG*, *OCT4* (*POU2F1*), and *KLF4* in tumor tuft cells (**Figure S2A**). Analysis of t-SNE data concurred with these findings and showed that *PALMD* expression was rare in normal colon epithelium, but expressed almost exclusively in CSCs and CRC tuft cells (**Figure 1G**), which represent two potential cellular origins of CRC^10,13^.

Given the proximity of *PALMD*+ CSCs and CRC tuft cells in the t-SNE projections, we sought to assess the lineage of these key populations in regards to *PALMD*. The HCCA scRNA-Seq data for CRC epithelial cells was filtered, dimensionally reduced, and projected using the UMAP technique, which demonstrated a smooth transition from CRC stem/TA-like epithelium (cE01) to CRC tuft cells (cE10)(**Fig S2B**). Projection of gene expression data onto the UMAP chart revealed a more discrete distribution for *PALMD* in CRC stem/TA-like epithelium compared to tuft marker *POU2F3* and CSC markers *LGR5* and *MYC* (**Fig S2C**). Next, we selected relevant cells for trajectory analysis (**Fig S2D**) and performed pseudotime analysis for tuft cells derived from stem cells (**Fig S3A**). Comparison of *PALMD* and tuft speciation marker *POU2F3* revealed a similar trajectory from CSCs to CRC tuft cells, while *LGR5* and *MYC* where primarily expressed in CSCs and not tuft cells (**Fig S3B**). Using the calculated values, we plotted the expression of *PALMD*, selected CSC markers (*LGR5*, *MYC*), tuft cell markers (*POU2F3*, *GNAT3*, *TRPM5*), and housekeeping gene β-actin (*ACTB*) as a function of pseudotime (**Fig S3C**). The plots revealed that *PALMD* shares dynamic expression similarities with both CSC and tuft cell markers and may be a factor in this cell lineage.

### PALMD knockdown impairs R-spondin2/LGR5/Wnt activation, cell growth, and stemness in CRC

Based on our pathological findings regarding *PALMD* expression, we selected undifferentiated cell lines HCT116 (CMS4), SW480 (CMS4), and DLD1 (CMS1) to assess the influence of *PALMD* expression on CRC growth and stemness^34^. To begin, we confirmed endogenous expression of *PALMD* and *LGR5* protein in all three cell lines (**Fig 2A**), and screened three shRNA constructs for their efficacy in knocking down *PALMD* in the HCT116 cell line (**Fig S4A**). Western blotting confirmed successful shRNA-mediated knockdown of PALMD with the first and third construct (shPALMD-1, shPALMD-3) in all cell lines, which led to notably decreased expression of CSC marker *LGR5* compared to controls (shCtrl)(**Fig 2B**). To understand how *PALMD* and *LGR5* might interact, we performed gene network analysis using the GeneMANIA algorithm, which predicted an interaction mediated by WNT-activator R-spondin2 (*RSPO2*) within the Lin *et al.* 2010 radiation hybrid genetic interactions dataset (**Fig 2C**). In agreement, analysis of TCGA’s PANCAN dataset confirmed a significant positive association between *PALMD* and *RSPO2* gene expression in CRC patient tumors (Pearson R = 0.46, Kendall’s tau = 0.314, P<0.0001)(**Fig 2D**). Given the central role of LGR/R-spondin signaling in the Wnt pathway, we sought to determine if *PALMD* is associated with its activation in CRC. Analysis of inferred pathway expression from TCGA PANCAN’s Paradigm dataset revealed strong activation of the Wnt signaling network in PALMD^High^ tumors (top 25^th^ percentile)(**Fig 2E**).

**Figure 2.**
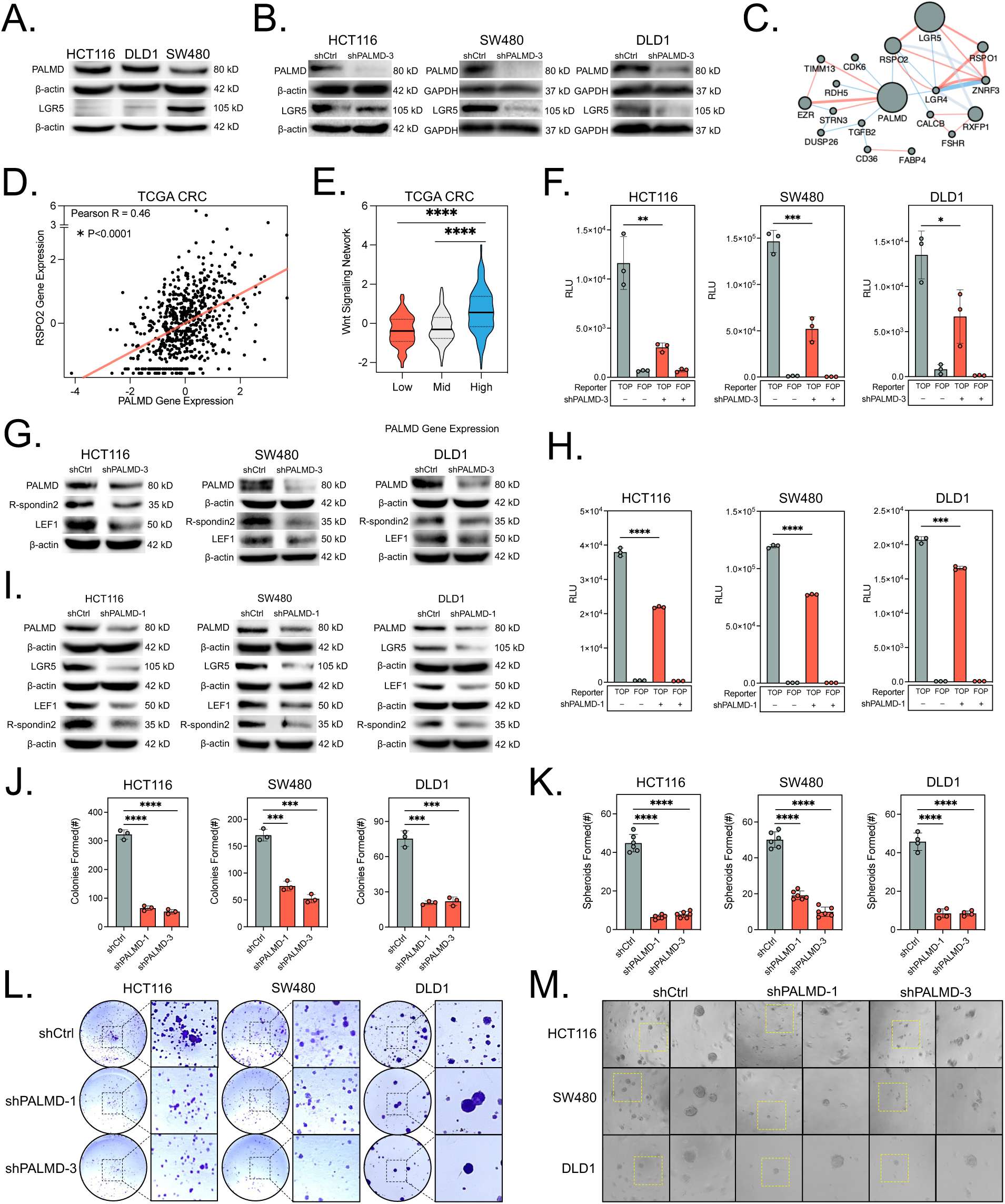
Silencing *PALMD* inhibits the R-spondin2/LGR5/Wnt signaling axis in colorectal cancer. **A.** Immunoblot showing baseline protein expression of *PALMD* and *LGR5* in HCT116, DLD1, and SW480 cells. **B.** Immunoblot demonstrating that shPALMD-3 transfection results in downregulation of LGR5 protein expression. **C.** GeneMANIA network analysis identifying a potential axis of interaction between *PALMD*, *RSPO2*, and *LGR5* as derived from the Lin *et al* radiation hybrid genetic interactions (2010) and Biogrid Small-Scale Studies datasets. **D.** Pearson analysis revealing a positive correlation between *RSPO2* and *PALMD* gene expression in CRC samples from TCGA’s Pan-Cancer Dataset. **E.** Comparison of WNT Signaling Network activation (z-score) compared to *PALMD* expression in CRC samples from TCGA’s Pan-Cancer PARADIGM pathway dataset, revealing significant activation of the WNT pathway in *PALMD* High tumors. **F.** Firefly luciferase reporter assay (TOPFlash) for TCF/LEF binding sites (TOP) and negative control (FOP) demonstrating statistically significant reporter suppression following shRNA-mediated knockdown of *PALMD* using shPALMD-3 in HCT116, SW480, and DLD1 cell lines. **G.** shRNA-mediated knockdown of *PALMD* using shPALMD-3 inhibits the expression of Wnt pathway agonist R-spondin2 and downstream Wnt transcription factor *LEF1*. **H.** TOPFlash assay demonstrating statistically significant reporter suppression following shRNA-mediated knockdown of *PALMD* using shPALMD-1 in HCT116, SW480, and DLD1 cell lines. **I.** Immunoblots demonstrating that shRNA-mediated knockdown of PALMD with shPALMD-1 results in decreased expression of WNT pathway members LGR5, LEF1, and R-spondin2. **J.** shRNA-mediated knockdown of *PALMD* (shPALMD-1/shPALMD-3) suppresses colony formation in HCT116, SW480, and DLD1 cell lines. **K.** shPALMD-1 and shPALMD-3 inhibit spheroid formation in HCT116, SW480, and DLD1 cell lines. **L.** Representative images of colony formation assays in HCT116, SW480, or DLD1 cells transfected with shCtrl, shPALMD-1, or shPALMD-3. **M.** Representative spheroids generated from shCtrl, shPALMD-1, or shPALMD-3 transfected HCT116, SW480, or DLD1 cells. *P<0.05, **P<0.01, ***P<0.001, ****P<0.0001.

To check directly for PALMD’s influence on Wnt pathway activation, we performed TCF/LEF-based binding assay (TOPFlash). In all CRC cell lines, shPALMD-3 led to a significant, 50% or greater reduction in TOPFlash reporter activation compared to shCtrl (p<0.05)(**Fig 2F**). To test the hypothesis that the expression of *PALMD* and R-spondin2 are correlated and involved together in Wnt pathway activation, we assessed the expression of R-spondin2 and downstream Wnt transcription factor *LEF1* after shPALMD-3 transfection in the cells, which have varying Wnt landscapes with DLD1 and SW480 harboring heterozygous and homozygous *APC* mutations respectively, and HCT116 harboring a single β-catenin mutant allele^35^. R-spondin2 and *LEF1* were downregulated by shPALMD-3 in all cell lines compared to controls (**Fig 2G**). To ensure the specificity of our findings, we repeated the Western blot and TOPFlash experiments with shPALMD-1. In confirmation, both TOPFlash (**Fig 2H**) and immunoblotting (**Fig 2I**) showed potent inhibition of reporter activation and *PALMD*, *LGR5*, *LEF1*, and R-spondin2 expression, respectively. To determine the functional effect of PALMD knockdown on cell growth and stemness, we used shPALMD-1 and shPALMD-3 and performed colony formation and spheroid assays. Both PALMD-targeted shRNAs resulted in a significant decrease in colonies formed and spheroid number in all 3 cell lines (**Fig 2J-M**). These *in vitro* findings implicate *PALMD* in a Wnt regulatory role with functional consequences to CRC tumorigenesis.

### PALMD promotes R-spondin2/LGR5/Wnt activation, β-catenin translocation, and CRC stemness

To assess whether *PALMD* can induce Wnt signaling through R-spondin2/LGR5, we used a PALMD^eGFP^ fusion construct (oePALMD), which was observed primarily in the cytoplasm of all 3 cell lines after transfection (**Fig S4B**). In all cell lines, oePALMD strongly promoted *LGR5*, *LEF1*, and R-spondin2 expression compared to oeCtrl as indicated by Western blot (**Fig 3A**). Furthermore, specific CSC marker *DCLK1* was significantly increased in oePALMD HCT116 and SW480 cells compared to oeCtrl, but no clear change was observed in DLD1 cells which expressed little to no *DCLK1* (**Fig S4C**). To further query Wnt-associated targets that may be related to *PALMD*, we performed RNA-Seq and PARADIGM integrated pathway level correlation analysis using CRC samples from TCGA’s PANCAN dataset. Significant positive associations were identified between *PALMD*, *RSPO2*, WNT receptors Frizzled-1 (*FZD1*) and Frizzled-8 (*FZD8*), WNT downstream transcription factor *LEF1*, and WNT/Frizzled complexes (WNT/LRP5-6/FZD), while negative correlations were found for WNT inhibitory E3 ubiquitin ligases *RNF43* and *ZNRF3* (**Fig 3B**).

**Figure 3.**
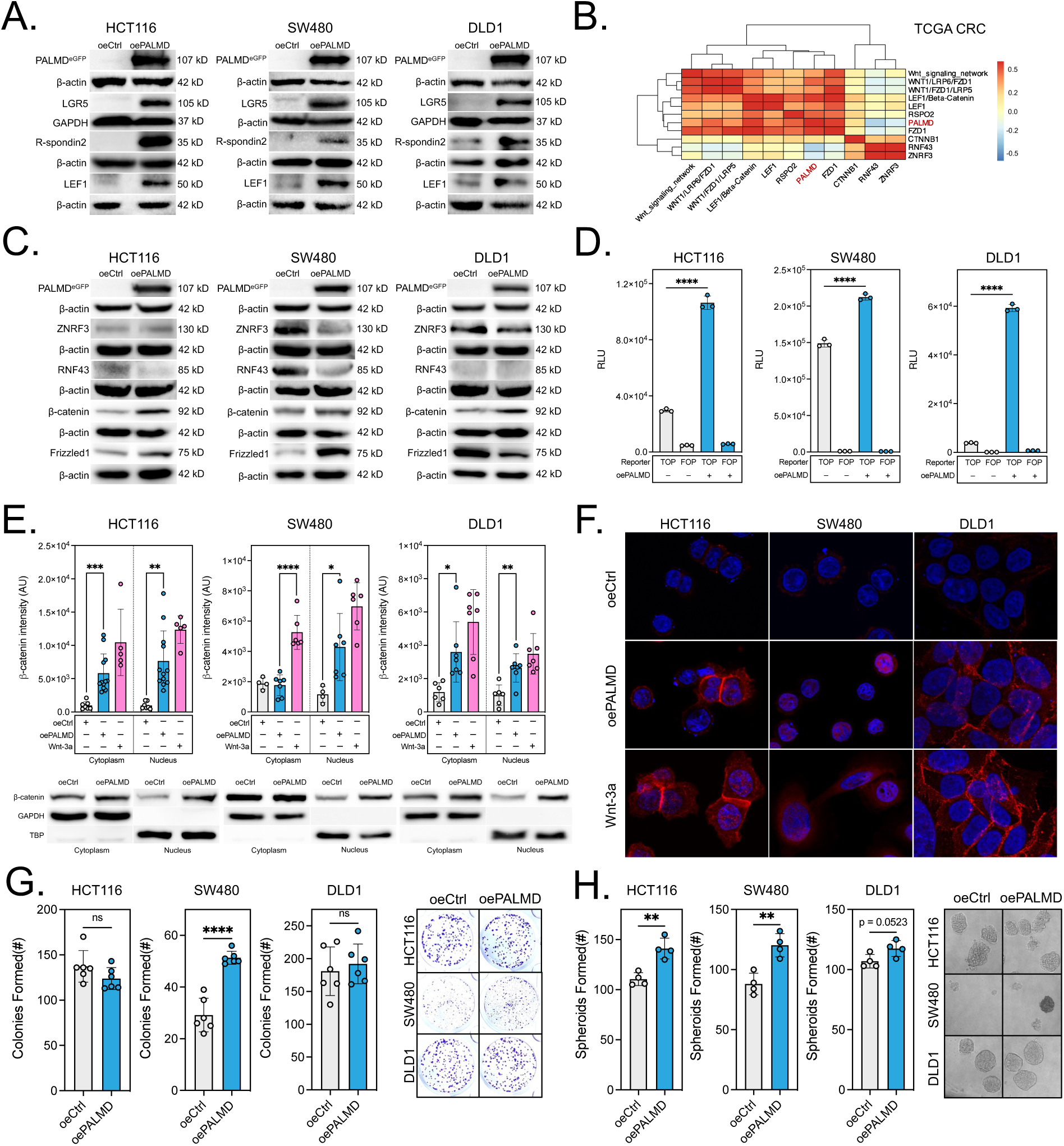
*PALMD* promotes CRC stemness, activates R-spondin2/LGR5/Wnt signaling, and induces nuclear translocation of β-catenin. **A.** Overexpression of *PALMD* (oePALMD) promotes expression of *LGR5*, *LEF1*, and R-spondin2 proteins in HCT116, SW480, and DLD1 cell lines. **B.** Gene co-expression analysis of PALMD, WNT/LRP/FZD complexes, and WNT-cascade genes using CRC samples from TCGA’s Pan-Cancer RNA-Seq (individual genes) and PARADIGM integrated pathway level (complexes) datasets. **C.** Immunoblotting of *PALMD* overexpression lysates from CRC cell lines demonstrating strongly reduced expression of Wnt inhibitor ubiquitin ligases *ZNRF3* and *RNF43* in SW480 and DLD1, and HCT116 and SW480 respectively; upregulation of Wnt transcription factor β-catenin in all 3 cell lines; upregulation of Wnt co-receptor Frizzled1 in HCT116 and SW480 cell lines. **D.** TOPFlash assay demonstrating significantly increased reporter activation following overexpression of *PALMD* in all 3 cell lines. **E.** Confocal immunofluorescence microscopy quantification demonstrating significant increases in nuclear β-catenin in HCT116, SW480, and DLD1 cells in oePALMD compared to oeCtrl (100 ng/ml exogenous Wnt-3a stimulation as positive control), and confirmatory cell fractionation immunoblots. **F.** Representative confocal microscopy of HCT116, SW480, and DLD1 cells after oeCtrl transfection, oePALMD tranfection, or exogenous Wnt-3a stimulation. **G.** Overexpression of *PALMD* promotes colony formation in the SW480 cell line. **H.** Overexpression of *PALMD* promotes spheroid formation in the HCT116 and SW480 cell lines. *P<0.05, **P<0.01, ***P<0.001, ****P<0.0001.

We sought to further strengthen our understanding of *PALMD*’s mechanism by investigating the expression of canonical Wnt pathway proteins in oePALMD cells. HCT116, SW480, and DLD1 oePALMD cells demonstrated increased expression of β-catenin, while HCT116 and SW480 oePALMD cells showed dramatically decreased expression of E3 ubiquitin ligase *RNF43* and increased expression of Wnt co-receptor Frizzled1 compared to oeCtrl. SW480 and DLD1 oePALMD cells showed a loss of E3 ubiquitin ligase *ZNRF3* compared to controls. However, DLD1 cells demonstrated no expression of *RNF43* and a decrease in the expression of Frizzled1 in oePALMD compared to oeCtrl (**Fig 3C**). Furthermore, increased TOPFlash reporter activation was noted in oePALMD compared to oeCtrl cell lines (P<0.0001)(**Fig 3D**).

To assess the influence of *PALMD* expression on downstream Wnt signaling, we performed immunofluorescence staining and nuclear fractionation to detect β-catenin in oePALMD compared to oeCtrl for all 3 cell lines. β-catenin expression was increased in all 3 oePALMD cell lines compared to oeCtrl, and showed more frequent nuclear localization in HCT116 and DLD1 cells which primarily express membrane/cytoplasmic β-catenin (**Fig 3E-F**). Given the central role of Wnt signaling in CRC growth and stemness, we performed colony formation and spheroid assays. oePALMD increased colonies formed in the SW480 cell line and spheroid number in both CMS4 cell lines (HCT116 and SW480). Although oePALMD DLD1 cells showed a modest increase in spheroid number, the result did not reach statistical significance (p=0.0523)(**Fig 3G-H**).

### Inhibition of β-catenin attenuates PALMD-mediated Wnt activation and stemness

CRC stemness and the transcription of key tumorigenic factors such as *LGR5* are reportedly regulated by Wnt/β-catenin signaling^36^. Given our findings in oePALMD cell lines, we sought to determine whether inhibiting downstream Wnt signaling through β-catenin could reverse PALMD’s Wnt-activating and stemness-inducing effect. We treated oeCtrl and oePALMD HCT116 or SW480 cells with 25 µM of β-catenin-TCF/LEF complex inhibitor iCRT3 or β-catenin degrader, MSAB for 16 h and performed western blotting analysis. iCRT3 was able to return *LGR5* and Frizzled1 expression levels to near oeCtrl baseline levels in oePALMD cells without significantly altering the expression of PALMD (**Fig 4A**). *PALMD*-induced β-catenin expression was also attenuated by MSAB treatment, which strongly reduced the expression of its previously reported downstream target, c-Myc^37^(**Fig 4B**). Next, we performed TOPFlash assay to determine if iCRT3 or MSAB inhibitor could reverse the effect of *PALMD*-mediated Wnt activation in HCT116 and SW480 cell lines. TOPFlash reporter activity was strongly reduced in both cell lines with both compounds as predicted (**Fig 4C-D**). To assess whether these findings were relevant to the functional effect of *PALMD* on cell growth and stemness, we repeated iCRT3 or MSAB treatment as described above and performed spheroid (HCT116 and SW480) assay. The number of spheroids formed from oePALMD cells after iCRT3 or MSAB treatment was dramatically reduced compared to vehicle (Veh) treated oePALMD cells (**Fig 4E-H**). Similar results were observed in the colony formation assay of iCRT3-treated SW480 cells (**Fig S4D**). Notably, in spheroid assays the extent of MSAB activity surpassed iCRT3, which may owe to MSAB’s direct β-catenin degradation mechanism. Combined, these findings support the notion that β-catenin inhibition can rescue CRC cells from tumorigenic PALMD-mediated Wnt activation and stemness *in vitro*.

**Figure 4.**
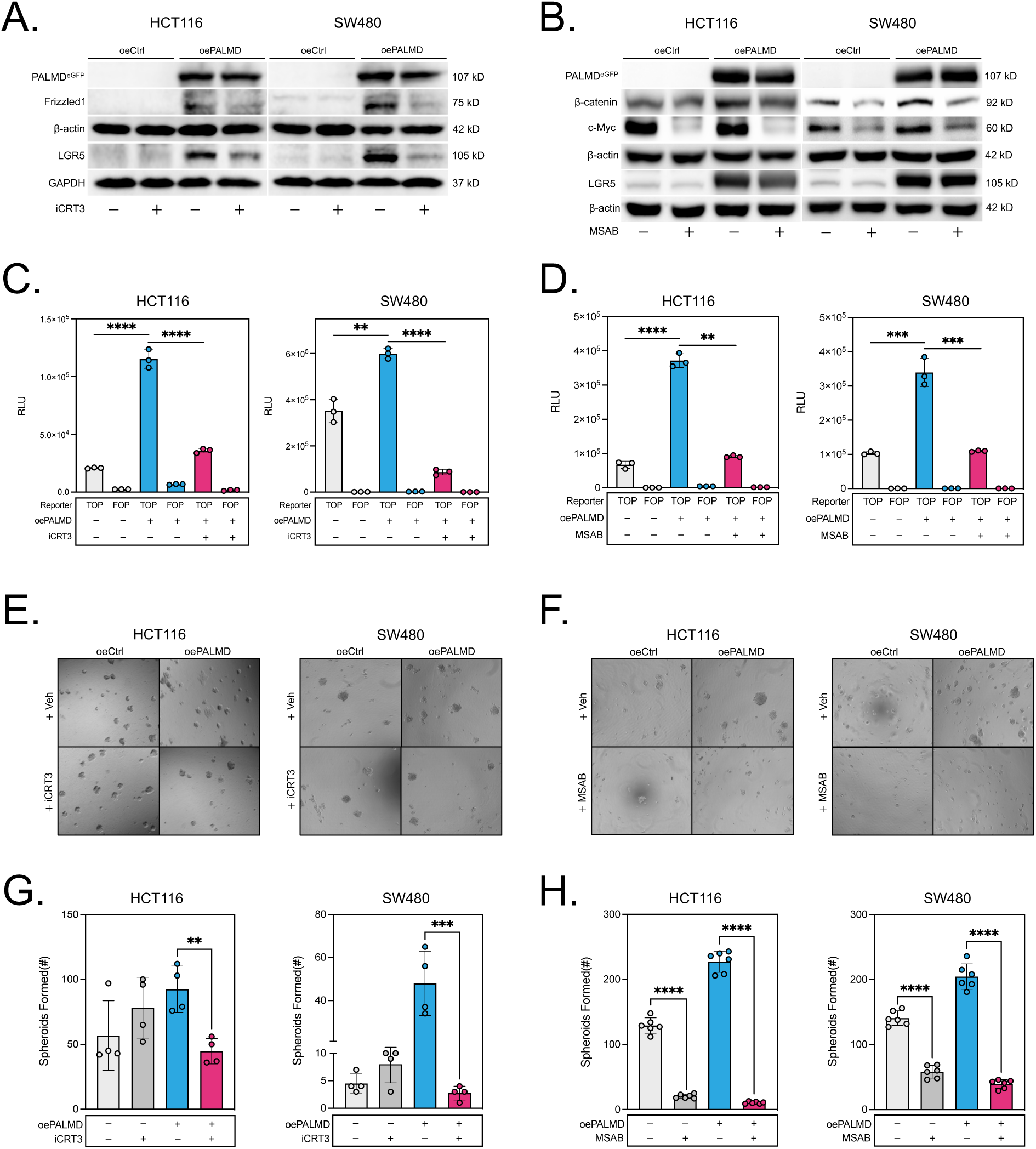
Inhibition of β-catenin reverses *PALMD-*induced WNT activation and stemness *in vitro*. **A.** Immunoblotting results demonstrating that PALMD-mediated expression of Wnt receptors Frizzled1 and *LGR5* are reversible by treatment with β-catenin/TCF-LEF complex inhibitor iCRT3 (25 µM). **B.** Immunoblotting results demonstrating that PALMD-mediated expression of β-catenin is attenuated by MSAB treatment (10 µM). **C.** TOPFlash assay demonstrating that *PALMD*-mediated TCF/LEF reporter activation is attenuated by treatment with iCRT3. **D.** TOPFlash assay demonstrating that *PALMD*-mediated TCF/LEF reporter activation is attenuated by treatment with MSAB. **E.** Terminal representative images of control (oeCtrl) and *PALMD*-overexpressing (oePALMD) HCT116 and SW480 spheroids treated with iCRT3. **F.** Terminal representative images of control (oeCtrl) and *PALMD*-overexpressing (oePALMD) HCT116 and SW480 spheroids treated with MSAB. **G.** Quantification of spheroid numbers in iCRT3-treated HCT116 and SW480 experiments revealing that *PALMD*-driven CRC spheroid formation is attenuated by iCRT3 treatment. **H.** Quantification of spheroid numbers in MSAB-treated HCT116 and SW480 experiments revealing that *PALMD*-driven CRC spheroid formation is attenuated by MSAB treatment. *P<0.05, **P<0.01, ***P<0.001, ****P<0.0001.

### PALMD activates the Wnt pathway by facilitating R-spondin2 secretion

To determine whether *PALMD*, R-spondin2, and *LGR5* interact on the protein level we performed co-immunoprecipitation using PALMD antibody and confirmed interaction in HCT116 and SW480 cell lysates (**Fig 5A**). Repetition of this assay using oeCtrl and oePALMD HCT116 and SW480 cell lysates demonstrated even greater accumulation of R-spondin2 with *PALMD* overexpression (**Fig S4E**). Given these findings, we sought to identify *PALMD’*s specific interaction partner using a bioluminescence resonance energy transfer (BRET2) assay. Co-transfection of RSPO2-Rluc8 and PALMD-GFP2 in HCT116 cells led to a significant increase in BRET ratio after 24 h, whereas LGR5-Rluc8 and PALMD-GFP2 co-transfection resulted in no apparent change (**Fig 5B**). Together these results lead us to conclude that *PALMD* interacts directly with R-spondin2.

**Figure 5.**
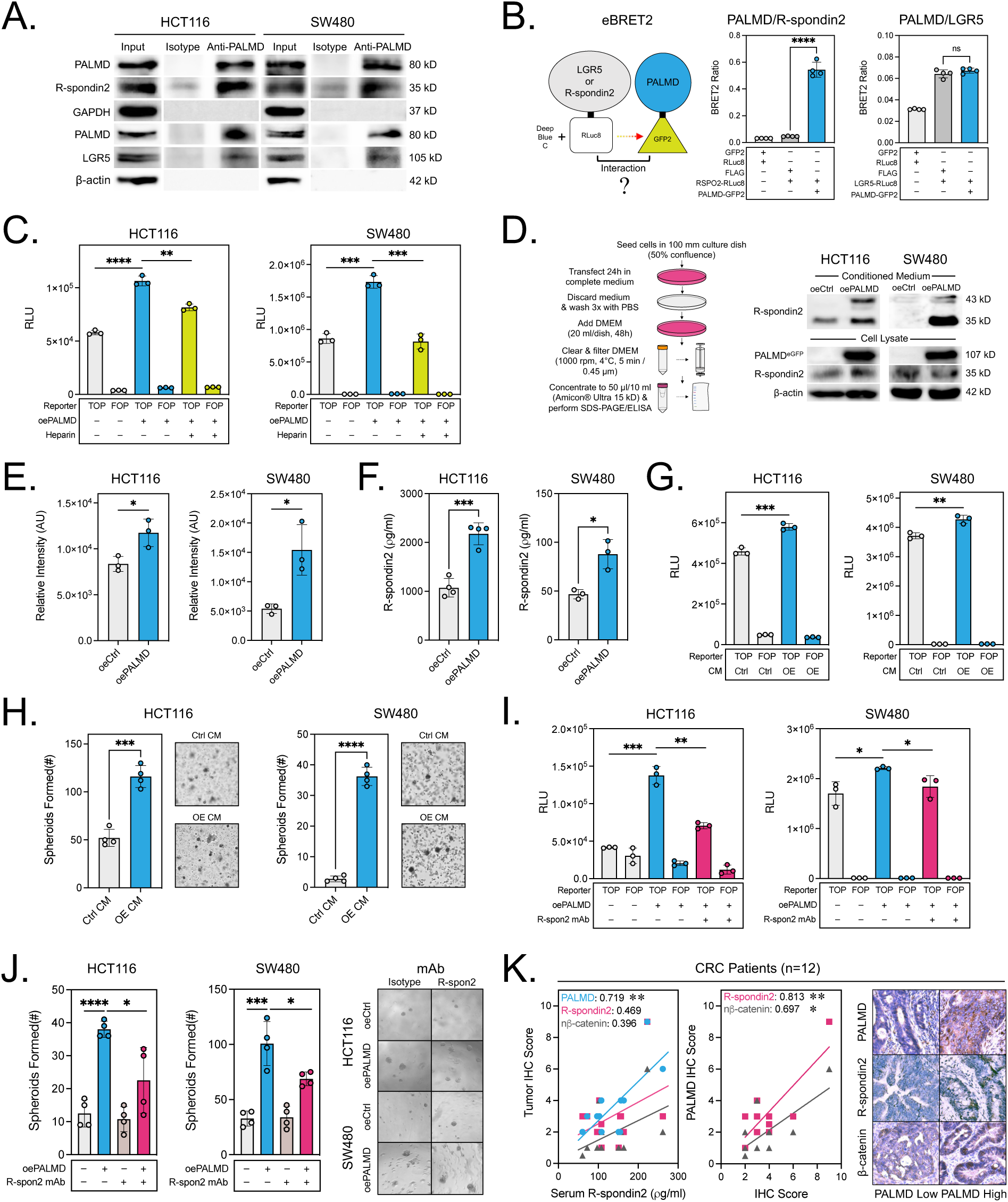
*PALMD* facilitates R-spondin2 secretion to promote paracrine Wnt activation and CRC stemness. **A.** Co-immunoprecipitation assay for anti-*PALMD* demonstrating that *PALMD* interacts with the R-spondin2/*LGR5* complex in HCT116 and SW480 CRC cell lines. **B.** Enhanced BRET2 assay results in co-transfected HCT116 cells demonstrating that *PALMD* specifically interacts with R-spondin2 but not LGR5. **C.** Heparin treatment significantly attenuates TOPFlash reporter activation in HCT116 and SW480 cells overexpressing *PALMD*. **D.** Immunoblotting demonstrating a strong increase in secreted R-spondin2 (35 kD) and it’s N-glycosylated form (43 kD) in conditioned medium from PALMD overexpressing HCT116 and SW480 cells compared to control, following 24 h transfection in antibiotic free DMEM complete medium and 48 h conditioning into serum-free DMEM. **E-F.** Quantitative Western blot and ELISA results demonstrating significantly increased R-spondin2 secretion in conditioned medium after *PALMD* overexpression in HCT116 and SW480 cells. **G.** TOPFlash assay demonstrating significant reporter activation in HCT116 and SW480 cells treated with conditioned medium from *PALMD* overexpressing cells for each cell line respectively. **H.** *PALMD* overexpressing cell-derived conditioned medium (OE CM) significantly promotes spheroid formation in HCT116 and SW480 cells compared to control medium (Ctrl CM). **I.** R-spondin2 blocking antibody (R-spon2 mAb) treatment significantly attenuates TOPFlash reporter activation in HCT116 cells overexpressing *PALMD*. I. Spheroid assay results demonstrating that R-spondin2 blocking antibody can inhibit *PALMD*-mediated stemness in HCT116 cells. **J.** Spheroid assay results demonstrating that R-spondin2 blocking antibody can inhibit *PALMD*-mediated stemness in SW480 cells. **K.** Pearson correlation analyses for matched serum R-spondin2 and tissue *PALMD*, R-spondin2, and β-catenin quantified by ELISA and immunohistochemistry respectively. *P<0.05, **P<0.01, ***P<0.001, ****P<0.0001.

R-spondins are secreted proteins that contain a thrombospondin domain and previous studies have demonstrated that soluble heparin can bind this domain and inhibit their ability to activate Wnt signaling^38–40^. To further interrogate the potential role of R-spondins in *PALMD*-mediated Wnt activation, we added heparin sodium to oePALMD HCT116 and SW480 cells for 48 h and performed TOPFlash assay revealing significant inhibition of Wnt reporter activity (**Fig 5C**). Similar findings were noted in oePALMD DLD1 cells (**Fig S4F**). Since R-spondin2 is a secreted protein, we sought to determine if *PALMD* regulates its secretion. Co-transfection of oeCtrl or oePALMD and RSPO2-RLuc8 plasmids in HCT116 and SW480 cell lines for 24 h revealed a significant increase in the culture medium luciferase activity of oePALMD/RSPO2-RLuc8 co-transfected cells compared to control, suggesting that PALMD may facilitate the secretion of R-spondin2 (**Fig S4G**). Next, we prepared oeCtrl and oePALMD conditioned medium (CM) by transfecting cells in complete medium for 24 h, followed by washing with PBS, and culturing in serum-free DMEM for 48 h. Immunoblotting of oePALMD CM produced from HCT116 and SW480 cells showed a potent increase in R-spondin2 expression compared to controls (**Fig 5D**). Additionally, a 43 kD form of R-spondin2 was also observed in the oePALMD CM lysate blots, which may represent an additional post-translationally modified form. We note that R-spondin2 expression was low in cell lysates obtained as controls for the CM western blot, showing slight upregulation in HCT116 and little change in SW480, suggesting that the majority of R-spondin2 has been matured and secreted into the CM at the experimental endpoint. Using Western blotting grey value data or ELISA to quantify R-spondin2 in HCT116 or SW480 oePALMD CM showed significant increases compared to oeCtrl CM (**Fig 4E-F**). These results demonstrate that *PALMD* facilitates R-spondin2 secretion in CRC cells.

Next, we sought to assess whether PALMD’s ability to activate Wnt signaling and stimulate functional stemness in CRC is related to its ability to promote the secretion of R-spondin2. TOPFlash assay with oePALMD CM (OE CM) showed a statistically significant increase in reporter activity compared to oeCtrl CM (Ctrl CM) in HCT116 and SW480 cells (**Fig 5G**). To test the functional implications of this finding, we cultured HCT116 or SW480 cells in Ctrl CM or OE CM and performed spheroid assay. Spheroid formation was significantly increased with OE CM exposure compared to controls in both cell lines (**Fig 5H**). To further confirm R-spondin2’s specific role in *PALMD*-mediated Wnt signaling and stemness, we employed R-spondin2 blocking antibody and performed TOPFlash and spheroid assays using HCT116 and SW480 cells. TOPFlash assay demonstrated that 0.25 µg/ml R-spondin2 blocking antibody (R-spon2 mAb), is able to significantly reverse *PALMD*-mediated Wnt activation (**Fig 5I**). Functionally, 0.25 µg/ml R-spon2 mAb treatment was able to inhibit *PALMD*-mediated spheroid formation, while having no effect on oeCtrl spheroid formation (**Fig 5J**). Finally, we sought to assess the relevance of these findings in CRC patients. Comparison of CRC patient serum R-spondin2 levels and PALMD expression by ELISA and immunohistochemistry respectively, revealed a significant positive correlation (Pearson r > 0.7). Moreover, immunohistochemistry-based quantification of R-spondin2 and nuclear β-catenin also revealed positive correlations with PALMD (Pearson r > 0.8 and 0.69, respectively) (**Fig 5K**). Together, these findings confirm that *PALMD* promotes Wnt activation and functional stemness in human CRC via stimulating paracrine R-spondin2 signaling, and that this process can be inhibited by R-spondin2 blockade.

### PALMD upregulates R-spondin2 and Wnt pathway and promotes CRC tumor growth in vivo

To investigate PALMD’s function *in vivo*, we established stable cell lines for SW480 oeCtrl and oePALMD, and confirmed their properties by Western blot and colony formation assay (**Fig S5A**). Additionally, we performed RNA-Seq analysis and found increased expression of CRC stem cell markers *LGR5*, *DCLK1*, *ASCL2*, *DACH1*, and *CD44* and pluripotency factors (*MYC*, *KLF4*, *SOX2*, *POU2F1*)(**Fig 6A**). Furthermore, *MYC* transcription factor targets where the top 2 categories in gene set enrichment analysis (**Fig S5B**). The stable cell line was implanted subcutaneously into athymic nude mice at 0.5×10^6^ cells per mouse at 5 weeks of age and tumor growth was monitored. Over the course of 53 days, oePALMD tumors showed a significantly increased rate of growth compared to oeCtrl (ANOVA P<0.0001)(**Fig 6B**). This finding was confirmed by measurement of excised tumor weight and volume (**Fig 6C-D**). Next we sought to determine the expression of *PALMD* and Wnt/CSC markers in tumor tissue by IHC and Western blot. IHC confirmed overexpression of *PALMD* and increased expression of R-spondin2 and *LGR5* in oePALMD compared to oeCtrl tumors (**Fig 6E**). Western blotting results confirmed overexpression of PALMD and increased expression of endogenous *PALMD*, R-spondin2, *LEF1*, *LGR5*, β-catenin, Frizzled1, and *DCLK1* (**Fig 6F**). Among these, densitometry-based quantification revealed statistically significant increases in endogenous *PALMD*, R-spondin2, and Frizzled1 (**Fig 6G**). Additionally, ELISA results for tumor lysates demonstrated an average R-spondin2 concentration of approximately 1.2 ng/µg compared to 0.55 ng/µg total protein for oePALMD and oeCtrl respectively (**Fig S5C**). Together these results confirm *PALMD* promotes Wnt pathway activation and tumor growth *in vivo*.

**Figure 6.**
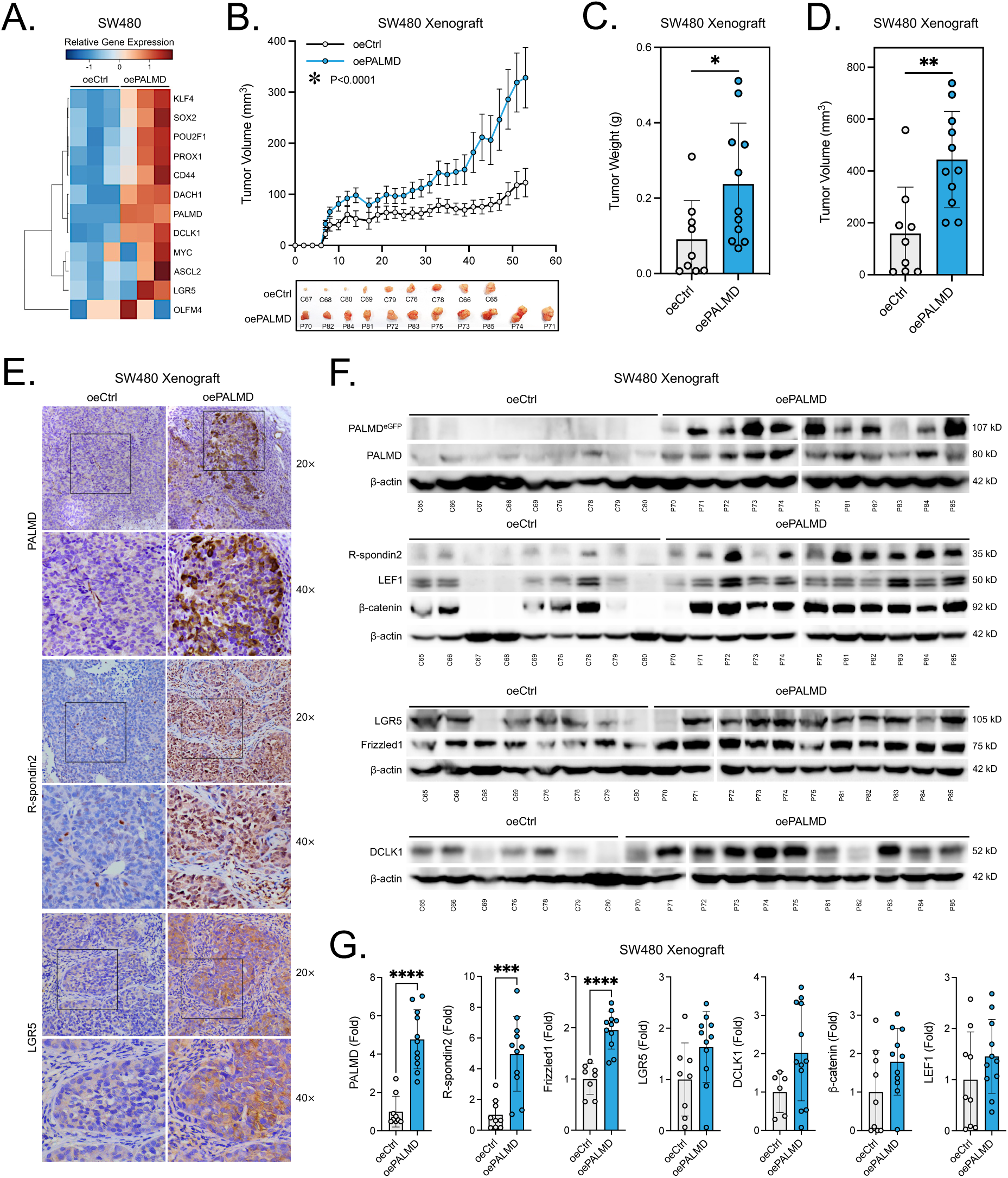
*PALMD* induces R-spondin2/Wnt pathway activation and promotes tumorigenesis *in vivo*. **A.** Heatmap demonstrating strong expression of previously reported CRC stem cell markers (*PROX1*, *CD44*, *DACH1*, *DCLK1*, *ASCL2*, *LGR5*) and pluripotency factors (*KLF4*, *SOX2*, *POU2F1*, MYC) in RNA sequencing data from SW480 cells stably overexpressing *PALMD* (oePALMD) compared to controls (oeCtrl). **B.** Xenograft growth curve and representative images showing a significant increase in tumor growth in SW480 oePALMD compared to oeCtrl tumors (ANOVA P<0.0001). **C.** The weight of SW480 oePALMD excised xenograft tumors was more than doubled on average compared to oeCtrl. **D.** The final excised tumor volume of SW480 oePALMD xenografts was more than doubled on average compared to oeCtrl. **E.** Immunohistochemistry staining confirming *PALMD* overexpression and demonstrating significantly increased R-spondin2 and *LGR5* expression in oePALMD compared to oeCtrl xenograft tissues. **F.** Immunoblotting showing: confirmation of PALMD-eGFP fusion protein overexpression and increased expression of endogenous *PALMD*, R-spondin2, *LEF1*, β-catenin*, LGR5*, Frizzled1, and *DCLK1* proteins in SW480 oePALMD compared to oeCtrl xenograft tissues. **G.** Immunoblot densitometry quantification demonstrating statistically significant increases in endogenous *PALMD*, R-spondin2, and Frizzled1 protein expression in SW480 oePALMD compared to oeCtrl xenograft tissues. *P<0.05, **P<0.01, ***P<0.001, ****P<0.0001.

### Emodin is a pharmacologic inhibitor of the PALMD/R-spondin2 axis

We next sought to identify pharmacologic inhibitors of *PALMD*. Literature analysis identified a single study, which reported downregulation of *PALMD* following treatment with 50 µM 6-methyl-1,3,8-trihydroxyanthraquinone (emodin) for 24 h in hepatocellular carcinoma cell lines Huh7, HepG2, and Hep3B^41^. To determine the relevance of this finding in CRC, we analyzed the proteomics and drug sensitivity dataset for emodin using the Cancer Dependency Map’s Cancer Therapeutics Response Portal (CTRP)^42–44^. Correlation analysis revealed a significant inverse association between PALMD protein expression and sensitivity to emodin (**Fig 7A**). Next, we performed Western blot for HCT116, SW480, and DLD1 cells treated with emodin for 48 h. *PALMD* and *LGR5* expression were significantly reduced in all 3 cell lines, and *DCLK1* expression was reduced in HCT116 and SW480 cell lines (**Fig 7B**), matching our findings with shPALMD and oePALMD (**Fig 2B**, **Fig 2I**, **Fig 3A**, **Fig S4C**). Immunofluorescence staining of SW480 and HCT15 cells, which express membrane *LGR5*, exhibited co-localization with *PALMD*. Treatment with emodin for 48 h significantly reduced the expression of both proteins (**Fig 7C**, **Fig S6A**). IHC staining of human CRC patient and SW480 oePALMD tumor xenograft tissue also suggested potential membrane localization for *PALMD* (**Fig S6B-C**). Finaly, Western blot analyses in HCT116, SW480, and DLD1 cells confirmed that emodin treatment can inhibit the expression of R-spondin2, Frizzled1, and β-catenin (**Fig 7D**).

**Figure 7.**
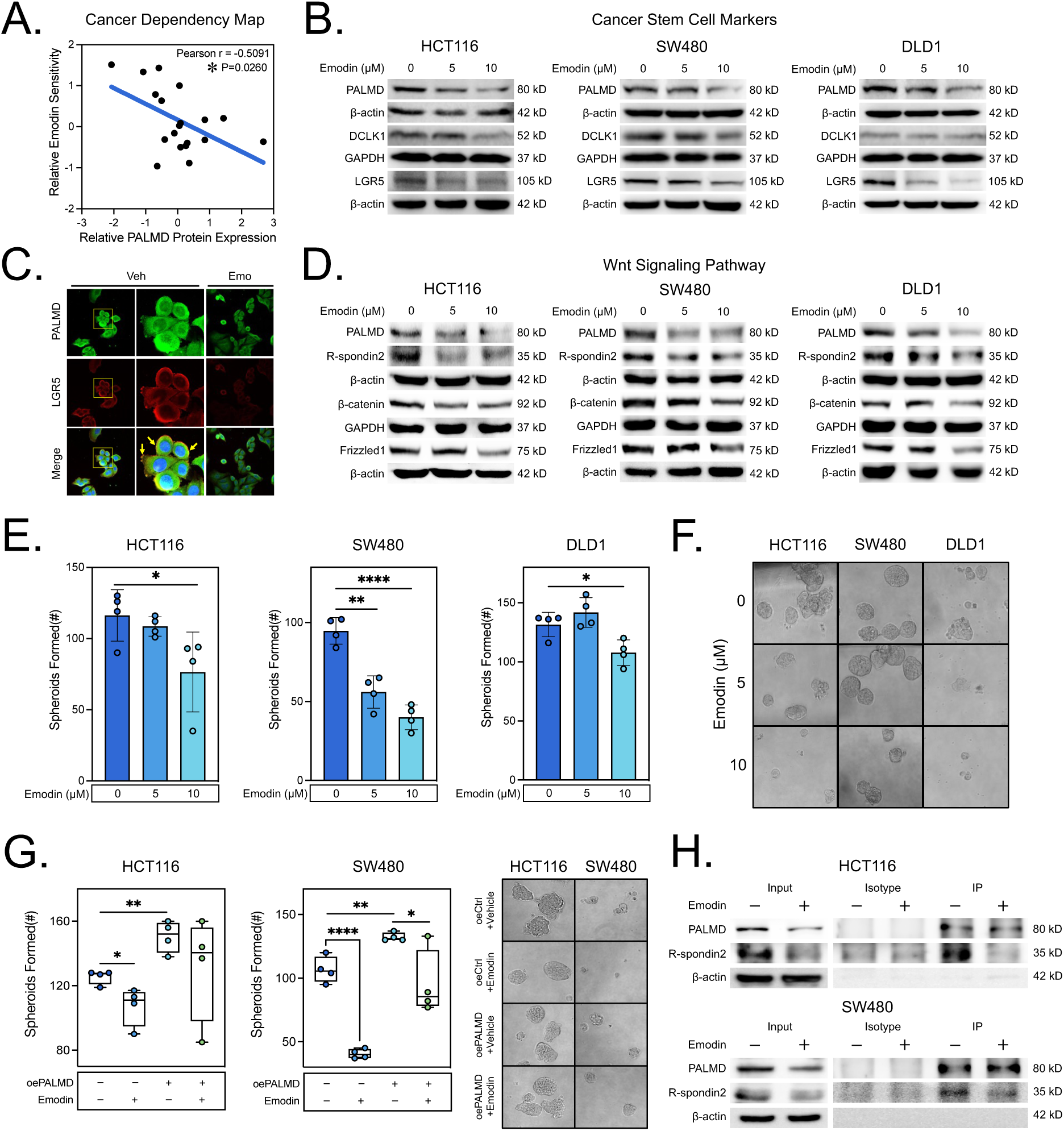
Emodin is a pharmacologic inhibitor of the *PALMD*/R-spondin2 axis. **A.** Correlation analysis of relative emodin sensitivity and *PALMD* protein expression in CRC cell lines as determined from the Cancer Dependency Map data (Pearson r = −0.5091). **B.** Immunoblotting in cell lines treated with 0, 5, or 10 µM of emodin for 48 h demonstrating consistent decreases in PALMD and LGR5 expression in HCT116, SW480, and DLD1 cell lines, and decreases in *DCLK1* expression in HCT116 and SW480 cell lines. **C.** Immunofluorescence staining for *PALMD* and *LGR5* in SW480 cells after 48 h of treatment with vehicle (Veh) or emodin (Emo; 10 µM) demonstrating *PALMD*/*LGR5* membrane co-localization and significantly decreased expression after treatment. **D.** Immunoblotting in cell lines treated with 0, 5, or 10 µM of emodin for 48 h demonstrating consistent decreases in *PALMD*, R-spondin2, β-catenin, and Frizzled1 expression in HCT116, SW480, and DLD1 cell lines. **E.** Quantification of number of spheroids formed after 0, 5, or 10 µM treatment with emodin, demonstrating significant dose-dependent decreases in SW480 and HCT116 spheroid formation and a significant decrease in DLD1 spheroid formation after 10 µM treatment. **F.** Representative images of spheroids formed from HCT116, SW480, and DLD1 cells treated with 0, 5, or 10 µM emodin. **G.** Quantification of number of spheroids formed after 0 or 10 µM treatment with emodin in oeCtrl or oePALMD lines, demonstrating *PALMD*-dependent reduction in emodin’s inhibitory effect on spheroid formation in HCT116 and SW480 cell lines. **H.** Co-immunoprecipitation results demonstrating that emodin treatment disrupts the interaction between *PALMD* and R-spondin2 in HCT116 and SW480 CRC cells. *P<0.05, **P<0.01, ***P<0.001, ****P<0.0001.

Next, we assessed the functional effects of emodin by proliferation, colony formation, and spheroid assay in all 3 CRC cell lines. Emodin showed limited anti-proliferative effects with IC_50_ values of 10.44, 19.69, and 26.67 µM in HCT116, SW480, and DLD1 cells respectively. These findings were confirmed by colony formation assay which showed no significant effect at 10 µM (**Fig S7A-B**). However, a significant inhibitory effect on spheroid growth was found in all 3 cell lines at 10 µM, suggesting that emodin has specific anti-stemness activity (**Fig 7E-F**). Next, we tested the effect of 10 µM emodin on spheroid formation from oePALMD HCT116 and SW480 cells compared to controls. Emodin was unable to significantly inhibit oePALMD HCT116 spheroid formation, and its effect on oePALMD SW480 spheroid formation was greatly diminished compared to oeCtrl (**Fig 7G**). To determine the activity of emodin in regards to the *PALMD/*R-spondin2 interaction, we performed co-immunoprecipitation in HCT116 and SW480 cells treated with emodin and observed a decrease in R-spondin2 from *PALMD* immunoprecipitates (**Fig 7H**). These findings indicate that emodin has specific anti-stemness activity in CRC, which is in part mediated by its inhibition of the *PALMD*/R-spondin2 axis.

### Inhibition of PALMD/R-spondin2 signaling attenuates CRC patient-derived organoid growth

To test the effect of *PALMD* in realistic models of CRC, we established patient-derived organoids (PDOs) from diagnostic biopsy samples. In total, 6 PDO models were established including 5 CRCs and 1 rectal adenoma (**Table I**, **Fig S8**). PDOs were screened for *PALMD* expression by Western blot, revealing variable expression (**Fig 8A**). Treatment of PDOs with emodin for 48 h resulted in decreased *PALMD* and R-spondin2 expression in 5/6 PDOs as determined by Western blot and IHC, with the exception of T0528, which had by far the highest expression level of *PALMD* (**Fig 8B**, **Fig S9**). After confirming the activity of emodin against *PALMD*/R-spondin2 in PDOs, we again treated PDOs with emodin for functional analysis. Through the course of treatment, a decrease in the number of PDOs was observed in all PDOs. However, the two highest expressers of *PALMD* (T1101, T0528) and the pre-cancerous rectal adenoma (T0519, p=0.072) did not reach statistical significance (**Fig 8C, F**). Staining for cell viability using calcein-AM and propidium iodide (PI) revealed a significant decrease in the ratio of living (calcein-AM) to dead (PI) cells in all models (**Fig 8D-E**). These results suggests that pharmacologic targeting of *PALMD* is a potentially effective therapeutic modality in CRC.

**Figure 8.**
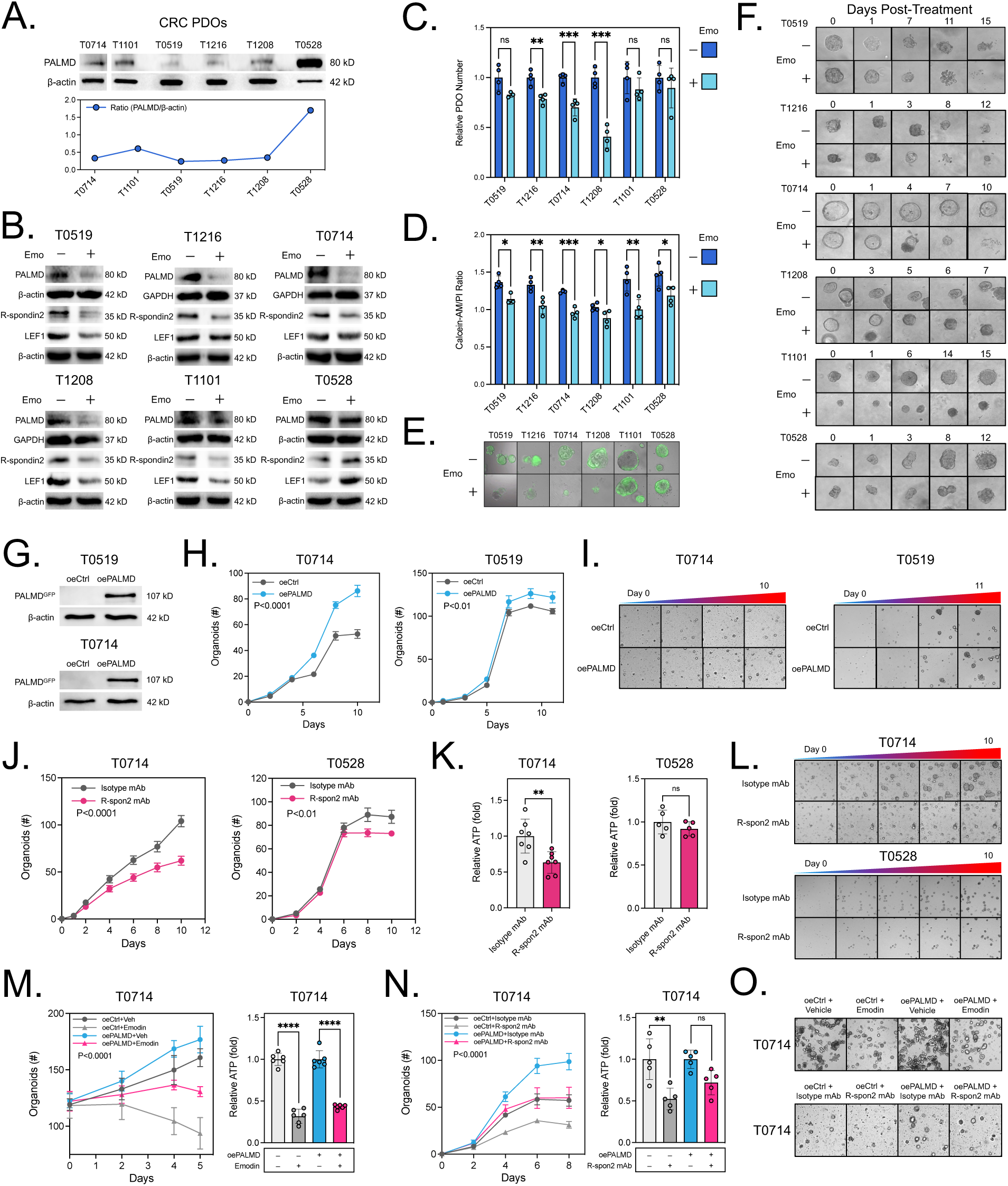
Pharmacological or physiological inhibition of the *PALMD*/R-spondin2 axis attenuates colorectal cancer patient-derived organoid (PDO) growth. **A.** Immunoblotting analysis of *PALM*D expression in 6 colonoscopy-derived PDO models demonstrating variable expression. **B.** Immunoblotting of CRC PDO lysates following treatment with 10 µM of emodin for 48 h demonstrating individualized reductions in *PALMD*, R-spondin2, and *LEF1* protein expression. **C.** Comparative analysis of relative PDO formation with or without 10 µM treatment with emodin revealing significant decreases in PDO number in CRC PDOs T1216, T0714, and T1208 but no significant difference in *PALMD* mid-high CRC PDOs T1101 and T0528 or rectal adenoma-derived PDO T0519. **D.** Calcein-AM/PI live/dead fluorescent cell staining quantification showing a decrease in PDO viability in all treated colorectal PDOs. **E.** Representative images of merged brightfield/calcein-AM live cell staining in PDOs treated with emodin. **F.** Representative images of individual PDO responses from each PDO model from day 0 to the final day of emodin treatment. **G.** Immunoblotting confirming overexpression of *PALMD* in T0519 and T0714 PDOs. **H.** Growth curves for T0519 and T0714 oeCtrl and oePALMD PDOs over the course of 7-11 days, demonstrating a statistically significant increase in PDO formation. **I.** Representative images of oeCtrl and oePALMD T0519 and T0714 PDOs throughout the course of the assay. **J.** Growth curves for T0714 (*PALMD* low) and T0528 (*PALMD* high) PDOs showing that 0.25 µg/ml R-spondin2 mAb treatment can inhibit PDO formation. **K.** ATP viability assay results showing that R-spondin2 mAb can significantly inhibit the viability of T0714 PDOs. **L.** Representative images of T0519 and T0714 after treatment with R-spondin2 mAb for 10 days. **M.** Growth curve/ATP viability results demonstrating that emodin (10 µM) can inhibit *PALMD-*mediated PDO formation. **N.** Growth curve/ATP viability results demonstrating that 0.25 µg/ml R-spondin2 mAb treatment can inhibit *PALMD-*mediated PDO formation. **O.** Terminal representative images of T0714 oeCtrl and oePALMD PDOs treated with emodin, R-spon2 mAb, or vehicle. *P<0.05, **P<0.01, ***P<0.001, ****P<0.0001.

**Table I.** Patient Characteristics of Tissue Donors for PDO Establishment.

| PATIENT ID | T0528 | T1101 | T1208 | T0714 | T1216 | T0519 |
| --- | --- | --- | --- | --- | --- | --- |
| **PATHOLOGY** | AdC | AdC | AdC | AdC | AdC | Adenoma |
| **LOCATION** | Sigmoid | Sigmoid | Sigmoid | Sigmoid | Rectum | Rectum |
| **AGE (YRS)** | 84 | 81 | 72 | 73 | 83 | 80 |
| **GENDER** | M | M | F | M | M | M |
| **CEA (NG/ML)** | 1.77 | 7.71 | 1.58 | N/A | N/A | 2.56 |
| **CA19-9 (U/ML)** | 2.1 | 18.31 | 6.5 | N/A | N/A | <2 |
| **AFP (NG/ML)** | 1.34 | 2.82 | 2.37 | N/A | N/A | 1.6 |

Abbreviations: Adc = Adenocarcinoma

To further ascertain the importance of *PALMD* in CRC PDO growth, we established stable *PALMD*-overexpressing PDOs using samples T0519 and T0714, which express relatively low endogenous levels of *PALMD*. Following confirmation of overexpression by Western blot (**Fig 8G**), we digested oeCtrl and oePALMD PDOs to single cells, and performed PDO establishment and growth assay. Both T0519 and T0714 oePALMD PDOs demonstrated a statistically significant increase in PDO formation compared to their respective oeCtrl counterparts (**Fig 8H-I**). Next, we sought to determine if R-spon2 mAb could inhibit PDO growth. Treatment of PDOs expressing low level (T0714) or high level (T0528) endogenous *PALMD* was performed, revealing that R-spon2 mAb inhibited growth in both PDOs as determined by the number of organoids formed (**Fig 8J, L**). However, a significant killing effect was only observed in *PALMD-*low T0714 as determined by ATP viability assay (**Fig 8K**). Next, we repeated R-spon2 mAb and emodin treatment in T0714 oePALMD organoids, and found that treatment abrogated enhanced *PALMD*-mediated PDO formation. Moreover, ATP viability assay revealed a significant decrease in oeCtrl viability after treatment with R-spon2 mAb, but no such effect was quantified in oePALMD PDOs (**Fig 8M-O**). These combined findings support *PALMD* as a novel and targetable factor in CRC stemness and tumorigenesis.

## Discussion

Our study shows that *PALMD* is a functional marker of CRC stem cells with an activator function in the Wnt pathway mediated by agonist R-spondin2. We initially identified *PALMD* using an ISC/CSC signature^33^ (**Fig 1A**, **S1A)**, and showed that its expression is interspersed in the crypt base epithelial stem cell region and expanded in CRC (**Fig 1B-C**, **S1B**), elevated in the 2 most aggressive CRC subtypes (CMS4/1)(**Fig 1D**), and predictive of CRC-specific survival and progression in those receiving first-line chemotherapy (**Fig 1E-F**, **S1E**). scRNA-Seq analysis further defined *PALMD*’s expression in a CRC CSC-tuft cell lineage (**Fig 1G-H**, **S2-3**), of which both cell types have been elegantly shown to act as cells-of-origin and continually fuel CRC growth and progression via *in vivo* lineage tracing and targeted cell ablation experiments in genetically engineered mouse models^10,13^. Given *PALMD*’s ability to strongly regulate *LGR5* expression (**Fig 2B**, **3A**) and the results of bioinformatics and network analyses (**Fig 2C-E**, **3B**), we tested *PALMD*’s effect on the Wnt pathway. *PALMD*-mediated Wnt activation was confirmed by Western blotting, TCF/LEF reporter assay (TOPFlash), and immunofluorescence of β-catenin nuclear translocation (**Fig 2F-I**; **Fig 3A, C-F**). Furthermore, through co-IP, BRET, immunoblotting, and ELISA assays, we determined that *PALMD* interacted with and promoted the secretion of Wnt agonist R-spondin2 (**Fig 5A-F**). Importantly, *PALMD*-mediated Wnt activation was attenuated by Wnt/β-catenin-inhibitors iCRT3 and MSAB, R-spondin-binding soluble heparin, and specific R-spondin2 blocking antibody – providing strong evidence for our proposed mechanism of action.

There are several potentially controversial aspects present in our study. First, in other cancer types studied to date, *PALMD* is not reported to induce a pro-cancerous phenotype. In uveal melanoma, *PALMD* expression was reported to promote migration and invasion^25^, and in breast cancer, its expression inhibited proliferation through the PI3K/AKT pathway^26^. In osteosarcoma, its expression promoted apoptosis after Adriamycin-induced DNA damage^24^. In our research, *PALMD* expression specifically induced functional stemness, while knocking down *PALMD* led to inhibition of this process in CRC. Here, we note that stemness is a functional property independent of others such as proliferation and migration/invasion. Indeed, the well-studied functional CSC marker *DCLK1* continues to generate controversy due to conflicting findings regarding its hypothetical potential (or at least the potential of some of its isoforms) to act as a tumor suppressor^45–49^. In further support of our findings, we provide clear evidence that *PALMD* potently activates the Wnt signaling pathway, which is known for its role in maintaining stemness. However, the R-spondin2 based mechanism which we demonstrate, may in itself generate controversy. Namely, although R-spondin2 has been shown to support tumorigenesis in multiple cancer types^30,50–52^, the evidence in CRC is varied. Wu *et al.* used *in vitro* and *in vivo* models to show that R-spondin2 can suppress Wnt activation and colony formation in CRC via interaction with *LGR5* which purportedly stabilizes the ubiquitin ligase *ZNRF3*. However, their findings demonstrated the opposite effect in CRC cell line HT-29^28^. Similarly, Dong *et al.* reported that R-spondin2 has tumor suppressor effects by activating non-canonical Wnt signaling in tandem with *Wnt-5a*, which is able to outcompete canonical Wnt signaling^53^. On the contrary, Zhang *et al.* showed that R-spondin2 promotes *LGR5* expression, epithelial-mesenchymal transition, spheroid formation, and invasion in CRC cells^54^. Furthermore, Chartier *et al.* demonstrated that treatment of CRC patient-derived xenograft models with specific R-spondin2 monoclonal antibody notedly inhibits tumor growth^29^. These findings and our own highlight the need for further studies to clarify R-spondin2’s role in CRC. In particular, we believe that this role may vary depending on CRC subtype. It is also likely that there are other yet to be discovered molecular players in the *PALMD*/R-spondin2 axis that promote the effects seen in our study.

## Methods

### Ethics Approval for use of Human and Murine Samples

Studies using human CRC tissue biopsies for PDO establishment were approved by the Ethics Committee of Fuzhou Traditional Chinese Medicine Hospital (FJTCM)(AF/SC-08/03.3). Studies of human CRC surgical tissue for IHC and ELISA were approved by the Medical Ethical Board of the Second Affiliated Hospital of Fujian University of Traditional Chinese Medicine (SPHFJP-T2022001-01). All patients provided informed consent for these studies, and the ethical and technical principles of biomedical research as described in the Declaration of Helsinki and the CIOMS international ethical guidelines were adhered to. After the patients signed the informed consent, the patient tissues and blood were collected during colonoscopy or surgical procedure. Each patient sample was pathologically confirmed. Clinical information can be found in **Supplementary Table I** and **Table I**. Mouse xenograft studies and wild-type C57/B6 tissue collection immunohistochemical studies were approved by the FJTCM IACUC #2021112 (application date: 09/09/2021).

### Cell Culture and Cell Line Generation

CRC cell lines (HCT116, DLD1, SW480, and HCT15) were purchased from the CAS cell bank, where their identity was confirmed. Routine PCR testing was carried out to ensure mycoplasma-free conditions. Cells were cultured in 10% FBS serum (Pan, ST30-3302) and 1% Antimycotic-Antibiotic. (Gibco, 15240062) in DMEM (Gibco, C11995500BT at 37°C in a 5% CO2-saturated humidity incubator. When the cell confluence reached 60%-70%, Cells were digested using 0.25% Trypsin-EDTA (Gibco, 25200056) and counted using a hemocytometer in preparation for experiments. For transfection, plasmid DNA for PALMD shRNA (shPALMD) or PALMD overexpression (oePALMD) and vector controls (shCtrl and oeCtrl) inserted into pcDNA3.1 were purchased from FH Biotechnology (Hunan, CN) and verified by Sanger sequencing. Cell lines HCT116, SW480, and DLD1 were transfected with plasmids according to the Lipofectamine 3000 (Thermo, L3000015) manufacturer instructions and confirmed by fluorescent microscopy (>50% transfection efficiency) and immunoblotting. To establish a stable cell line, oeCtrl and oePALMD SW480 cells were seeded at 10 cells per well into a 96 well plate, allowed to attach, and selected with G418. This process was repeated 3 times in both cell lines until integration was confirmed by Western blotting after G418 removal and passaging.

### CCK8 Assay

Cells were seeded into 96-well plates (100μl/well) at 1×10^5^/ml. When confluency reached 50%, they were treated with varying concentrations of emodin (MCE, HY-14393) from 0 (DMSO only) to 250 µM. After 72 h, 10 μL/well of CCK-8 reagent (Apex, K1018-100) was added and the cells were placed into an incubator for 2 h. Finally, the absorbance value was detected at a wavelength of 450 nm using a multifunctional microplate reader.

### Colony Formation Assay

Cells were seeded in 24-well plates at 0.3×10^3^/ml cells and allowed to attach for several hours. Next, cells were cultured and, when relevant, treated with vehicle (DMSO), emodin, or iCRT3 (25 µM; MCE, HY-103705) for 7-10 days. At termination, the medium was removed, wells were washed with PBS, and cells were fixed with 10% neutral buffered formalin (NBF; Solarbio, G2161) for 15 min at room temperature. After fixation, the wells were washed again and stained with 0.1% crystal violet (Solarbio, G1062) for 10 min. Crystal violet-stained cells were then washed gently with water several times. A colony was considered to contain 50 cells or more when counted under the microscope.

### Spheroid Formation Assay

Cells (0.5×10^3^/25 μl) were mixed gently with 25 μl of growth factor-reduced Matrigel (Corning, 356231) per well and then seeded into a preheated 96-well plate, which was placed in an incubator for 30 min for matrigel solidification. 100 μL/well of DMEM containing 0.5% FBS was added to each well and, when relevant, treated with vehicle/control (DMSO or isotype monoclonal antibody, MCE, HYP99979), emodin (0, 5, or 10 µM), iCRT3 (25 µM), or R-spondin2 blocking monoclonal antibody (RND, MAB3266). For conditioned medium (CM) assays, treatment consisted of oeCtrl or oePALMD-derived concentrated CM with pen/strep and containing no FBS. During the experiment, medium was replaced every 3 days. After 7-10 days, the plate was washed with PBS and 200 µl/well of NBF was added for fixation at room temperature for 30 min. Each spheroid was counted and pictures (20X) were taken under a microscope. Spheroids were considered to have a diameter value of 100 μm or greater.

### Immunocytochemistry

Cells were diluted to 1×10^5^ cells/ml and seeded in a u-Slide 8 well plate (Ibidi, 80806) at a volume of 200 µl/well. After adherence, they were treated with 10 µM emodin or DMSO for 48 h, if appropriate. Fixation was performed with 10% NBF for 15 min followed by washing with PBS. Permeabilization was performed using 0.1% Triton-X for 15 min, followed by blocking with 10% donkey serum (Solarbio, SL050) for 30 min at room temperature. Afterwards, wells were washed with PBS, and primary antibody was added overnight at 4°C. Primary antibodies used included: PALMD (Proteintech, 16531-1-AP, 1:100) and LGR5 (Abcam, ab273092, 1:100). The next day, primary antibody was removed and wells were washed with PBS. Secondary antibodies were then incubated for 1 h at room temperature in the dark. Secondary antibodies used included: donkey anti-rabbit IgG (Abcam, ab150073, 1:500) and donkey anti-mouse IgG (Abcam, ab150108, 1:500). After incubation, secondary antibodies were removed and wells were washed with PBS. 300 µl/well of DAPI (Beyotime, 1005) was then added and incubated in the dark according to manufacturer instructions. Immunofluorescence images were taken using a Leica DMI4000 B fluorescent microscope (20X) and the resulting photographs were merged using ImageJ.

### Confocal Microscopy and Quantification

For confocal microscopy, glass coverslips were sterilized by UV light for 30 min and then seeded with HCT116, SW480, or DLD1 cells. After cells attached to the coverslips overnight, transfection was performed with oeCtrl or oePALMD plasmid for a total of 24 h. As a positive control, mock transfected cells were cultured under the same conditions for 22 h and then stimulated with 100 ng/ml Wnt-3a (MCE, HY-P70453B) for 2 h. Afterwards, the coverslips were washed with PBS, fixed with 10% neutral buffered formalin for 15 min, washed again with PBS, permeabilized with 0.1% Triton-X 100, washed with PBS, and then blocked in 1% BSA for 2 h. Following blocking, the coverslips were washed with PBS and then incubated with β-catenin primary antibody (Abcam, ab32572, 1:250) overnight at 4°C. The next day, the antibody was collected and the coverslips were washed 3 times with PBS. AlexaFluor-647 secondary antibody (Proteintech, SA00014-9, 1:100) was then added for 2 h at room temperature. The coverslips were counterstained with DAPI and mounted onto glass slides for imaging under a Stellaris 8 confocal microscope (Leica). To quantify confocal microscopy results, a custom pipeline was performed in CellProfiler v4.2.7 (BROAD institute). For quantification of nuclear β-catenin, DAPI (blue) and β-catenin (red) channel images were imported and converted from RGB to gray using the *ColorToGrey* function. Next, nuclei were positively identified using *IdentifyPrimaryObjects* (300 – 10000 pixels and discard edge objects), and the result objects were input to *MaskImage*. Finally, the masked region was selected and used to measure the β-catenin channel nuclear intensity using *MeasureObjectIntensity*. For quantification of cytoplasmic β-catenin, *ColorToGrey* and *IdentifyPrimaryObjects* were run as described above. Next, *IdentifySecondaryObjects* was performed using the ‘Watershed – Gradient’ method, ‘Adaptive’ threshold, and ‘Otsu’ thresholding method. The *IdentifyPrimaryObjects* output (nucleus) was subtracted from the *IdentifySecondaryObjects* output (cytoplasm) using the *IdentifyTertiaryObjects* function. Finally, *MeasureObjectIntensity* was used to obtain the intensity of β-catenin in the cytoplasm.

### Protein Extraction

Cells were seeded into 6 cm cell culture dishes at 1.0×10^6^/ml and allowed to reach 50% confluence. After culturing and treatment, the medium was aspirated and the plate was washed with PBS. 200 μL of cell lysis buffer containing mammalian protein extraction reagent (MPER; Pierce, 78501), PMSF (Keygen, KGP610), and protease inhibitor cocktail (MCE, HY-K0010) was added to each well. Following 1 min of incubation, a cell scraper (Greiner, 541070) was used to collect the lysate into a 1.5 ml tube and the lysate was centrifuged (4°C, 10000 rpm, 10 min). The total protein concentration of the supernatant was then quantified by BCA assay. The remaining samples were denatured in a heat block using 5x SDS dye at 100°C for 5 min and stored at −20 °C. Similarly, for xenograft tumor tissue, tissue wash washed 3 times with PBS and then protein was extracted into the lysis buffer described above using a tissue grinder.

### Preparation of Conditioned Medium

Cells were cultured in 2 × 100 mm^3^ cell culture dishes per transfection group to 70% confluence. Cells were transfected with oeCtrl or oePALMD plasmid using lipofectamine 3000 transfection reagent as described above. After 24 h, complete medium containing transfection reagent was removed, successful transfection was confirmed by microscopy, and dishes were washed gently 3 times with sterile PBS. 25 ml of DMEM containing no FBS or antibiotics was then added to each plate and allowed to culture for 48 h. After 48 h, CM was collected, centrifuged at 1000 rpm at 4°C for 5 min and filtered through a 0.22 µm PES mesh strainer to clear any detached cells, and concentrated using Amicon Ultra-15 protein concentrators (Merck, UFC901008) with a 10 kD molecular weight cutoff to equal volume per group (250 µl). 50 µl of CM was combined with 50 µl of MPER containing 1x PMSF and protease inhibitor cocktail and then denatured on a heatblock with 5x SDS dye. The remaining CM was used freshly for TOPFlash reporter and spheroid assays and additional CM was stored at −80°C for subsequent use within 1 week.

### Generation of Colorectal Cancer Patient-Derived Organoids

CRC patient colonoscopy tissue were added to 15 ml centrifuge tubes containing cold D-PBS (STEMCELL, 37350) with Primocin (Invivogen, ant-pm-2) and stored at 4°C for transportation. Samples were washed 10 times with 10 ml of cold D-PBS in a 15 mL centrifuge tube. The tissue was then transferred to a 1.5 ml tube using a 1 ml sterile tip, cut into small pieces (about 5 mm) using sterile surgical scissors, resuspended in 1 ml of Gentle Cell Dissociation Reagent (GDCR, STEMCELL, 07174), and transferred to a new 15 mL tube. The 1.5 mL tube was washed with 1 ml of GCDR, which was then transferred to the same 15 ml tube. Additional GCDR reagent was added to the 15 ml tube to 10 ml total volume and the tube was placed on a shaker at 40 rpm at 37°C for 1 h. Afterwards, the tube was centrifuged (290g, 4°C, 5 min) and the supernatant was discarded. 1 ml of cold DMEM/F12 with 15 mM HEPES medium (STEMCELL, 36254) containing 1% BSA (Solarbio, A8010) was added and the pellet was pipetted up and down 15 times vigorously and transferred to a new 15 ml tube through a 70 μm Reversible Strainer (STEMCELL, 27216). This step was repeated to ensure all crypts were obtained from the 15 ml tube. Following filtration, the new tube was centrifuged (200g, 4°C, 5 min) and excess liquid removed. Crypts were then seeded into a 50 µl 50% matrigel dome (2000 crypts/well) in a preheated 24-well plate. After solidification in a 37°C incubator, 750 µl/well of complete IntestiCult^TM^ Organoid Growth Medium (OGM; STEMCELL, 06010) and 10 µM Y-27632 (MCE, HY-10583) was added to each well. OGM was replaced every 2 days. Organoid establishment was observed and photographs were taken to record growth until maturity. After maturation, organoids were passaged for cryopreservation and immediately used in experiments.

### Lentiviral infection of CRC PDOs

Lentivirus for oeCtrl and oePALMD were purchased from GeneChem and expressed from the GV492 vector. Overexpression was confirmed by qPCR (GeneChem) and by Western blot analysis with specific antibody. To establish stable oePALMD and oeCtrl PDOs, healthy PDOs were digested with TrypLE (Thermo, 12604013), filtered through a 70 μm strainer, seeded at 2 × 10^5^ cells/well in a 24 well plate in OGM containing 10 µM Y-27632 and 7.5 × 10^6^ TU of relevant lentivirus, and incubated at 37°C overnight. On the second day, adherent PDO cells were detached using TrypLE and cells were seeded into matrigel domes at 1 × 10^5^ cells per well as described above. After matrigel solidification, cells were fed with OGM containing 10 µM Y-27632 and 1µg/ml puromycin dihydrochloride (MCE, HY-B1743A). PDOs were monitored for selection by fluorescent microscopy with continuous changes of puromycin-containing OGM until reaching 100% positivity. Subsequently, Western blotting was performed to confirm overexpression.

### CRC PDO Growth Assays

For assessment of oePALMD and oeCtrl PDO growth, PDOs were digested to single cells with TrypLE. A 96 well plate was seeded at 200 cells/well in 6 µl of 50% matrigel with OGM and baseline images were taken with a MuviCyte (Revvity). Every 2 days, OGM was changed, images were taken, and PDOs were counted. For assessment of emodin activity, PDOs were seeded at 200 crypts/well in a pre-warmed black transparent bottom 96-well plate. Following maturation, organoids were counted under a microscope and pictures were taken (20x) to establish baseline values. The medium was replaced again before treatment with 10 µM emodin or equivalent vehicle. During the experiment, the medium was replaced every day and photographs were taken every 1-2 days as necessary. The experiment was continued 7-15 days depending on organoid health and growth in the vehicle control group, which varies by individual PDO. At termination, Calcein-AM (Keygen, KGMP012-1) with a final concentration of 5 µM was added for 1 h at 37°C. After washing away the excess Calcein-AM, propidium iodide (Beyotime, ST1569) was added at a final concentration of 1 µg/ml and incubated 15 min at room temperature. The relative fluorescence value was detected using a multiplate reader and representative photographs were taken using a Leica DMI4000 B fluorescent microscope (20X).

### Extraction of Proteins from CRC-PDOs

PDOs were seeded at 2000 crypts/well in a preheated 24-well cell culture plate and cultured as described above. For emodin experiments, fresh medium containing 10 µM of emodin or vehicle was replaced every day for a total of 72 h. After 72 h, the medium was removed and the plate was washed 3 times with D-PBS. Another 1 ml/well of D-PBS was added to the plate and a pipette was used to scrape and pipet up and down 10 times to destroy the matrigel structure. The resulting liquid was transferred to a 15 ml tube and centrifuged (4°C, 290g, 5 min). Following centrifugation, the supernatant was discarded and 200 µl of protein lysis buffer was added to each tube to fully resuspend the pellet, which was then transferred to a 1.5 ml tube and incubated on a shaker for 5 min (4°C, 40 rpm). The samples were resuspended again 10-15 times and centrifuged (4°C, 10000 rpm, 10 min). After centrifugation, the total proteins in the supernatant were quantified and denatured as described previously.

### Western Blot Assay

Protein lysates were denatured on a 100°C metal heat block for 5 min, loaded onto an SDS-PAGE gel in equal amounts according to BCA assay results, separated by electrophoresis, and transferred onto a PVDF membrane. After transfer, the membrane was blocked in 1X blocking buffer (Abcam, ab126587) for 1 h and incubated with primary antibody at 4°C overnight. The primary antibodies used include: PALMD (Proteintech, 16531-1-AP), LGR5 (Bioss, bs-20747R), DCLK1 (Abcam, ab109029), R-spondin2 (Bioss, bs18876r), LEF1 (CST, C12A5, 2230), ZNRF3 (Bioss, bs-7007R), RNF43 (Bioss, bs-9141R), β-catenin (Abcam, ab32572), Frizzled-1 (Immunoway, YT773), ATP1A1 (Na+/K+ ATPase 1a; Proteintech, 14418-1-AP), GAPDH (Proteintech, 60004-1-lg), and β-actin (Proteintech, 66009-1-lg). On the second day, the membrane was washed three times with TBST for 5 min each time and then incubated with horseradish peroxidase-conjugated, species-specific secondary antibody (Proteintech) for 2 h. The results were imaged using a Bio-Rad Gel Doc XR+ or a Cytiva ImageQuant™ 800 chemiluminescence system.

### Co-Immunoprecipitation

For co-immunoprecipitation (co-IP), 2 × 10 cm dishes of cells were cultured and allowed to reach a confluence of 90%. Protein was collected in MPER containing 1 x PMSF and protease inhibitor cocktail and then incubated with 11 µg of PALMD antibody (Proteintech, 16531-1-AP) or rabbit IgG isotype control (Sino Biological, CR1) on a rotating platform overnight at 4°C. Next, Protein A/G beads (Beyotime, P2108), were washed 3 times with TBST, and then the antibody-protein complexes were incubated with the beads on the rotating platform for 1 h at room temperature. A magnetic stand was used to capture the complexed Protein A/G beads, which were then washed 3 times with TBST, and eluted into 1x SDS for Western blot.

### R-spondin2 ELISAs

For R-spondin2 ELISA assay from xenograft tissues, extracted protein from SW480 oeCtrl and oePALMD tumors were collected. BCA assay was performed prior to ELISA and sample loading amounts were adjusted accordingly. The ELISA was carried out as described in the manufacturer instructions (CUSABIO, CSB-EL020551HU). For conditioned medium and human CRC patient serum, a high-sensitivity human R-spondin2 ELISA kit was used according to the manufacturer instructions (Signalway Antibody, EK7505). ELISA results were calculated using the GainData® (Arigo biolaboratories) software.

### TOPFlash Reporter Assay

TOPFlash (Wnt reporter) and FOPFlash (control) plasmids were purchased from Beyotime (D-2505 and D-2507 respectively). Cells were co-transfected with TOPFlash, FOPFlash, and target plasmids (oeCtrl, oePALMD, shCtrl, shPALMD) as required according to the lipofectamine 3000 manufacturer instructions. For untreated oePALMD experiments, transfection was performed for 24 h. For untreated shPALMD experiments, transfection was performed for 72 h. For iCRT3, drug or vehicle (DMSO), cells were washed with PBS 8 h post-transfection before drug addition in new complete medium and cells were cultured until 24 h post-transfection. For heparin sodium (STEMCELL, 07980; 100 µg/ml) and R-spondin blocking antibody (0.25 µg/ml), cells were washed in PBS 8 h post-transfection and drug or vehicle (PBS and isotype control antibody respectively) were added in new complete medium until 48 h post-transfection. Following transfection/treatment, cells were again washed with PBS, lysed, centrifuged to clear cell debris, and then lysis supernatant was mixed 1:1 with luminescence detection reagent (Beyotime, RG005) in a white 96 well plate while avoiding light exposure. Relative luminescence units (RLU) were detected on a Tecan Spark multifunctional plate reader.

### Bioluminescence Resonance Energy Transfer (BRET) Assay

Enhanced BRET2 (eBRET2) assay was performed using co-transfection of pcDNA3.1 plasmid pairs: RLuc8-Vector and GFP2-Vector, LGR5-RLuc8 and FLAG (empty), RSPO2-RLuc8 and FLAG, PALMD-GFP2 and FLAG, LGR5-RLuc8 and PALMD-GFP2, RSPO2-RLuc8 and PALMD-GFP2. After 24 h of transfection, 1.8 × 10^5^ cells/well were seeded into a sterile, opaque white plate (n=4). Measurements were taken on a Berthold Tristar 5 Multimode Reader after the addition of 5 µM of coelenterazine 400a in the LightCompass® software using both BRET2 channels, and the acceptor/donor ratio was analyzed to determine the extent of energy transfer.

### Immunohistochemistry, Immunofluoresence, and Pathological Analysis of Stained Tissue

Formalin fixed paraffin embedded (FFPE) tissue sections were deparaffinized and subjected to antigen retrieval with EDTA (human tissue) or citrate (mouse tissue and PDOs) buffer, followed by endogenous peroxidase blocking (Immunoway, RS0053) for 30 minutes to avoid nonspecific binding. Following blocking, primary antibody was added onto the tissue slides at 4°C overnight with a specific and verified primary antibody against PALMD (1:1000, Sigma, HPA030549), β-catenin (1:250, Abcam, ab32572), or R-spondin2 (1:300, Bioss, bs18876r). The sections were washed 3 times with 1x PBST solution for 5 min and then incubated with secondary antibody for 30 min at room temperature. After a second round of washing, the sections were stained with DAB (Origene, ZLI-9018) and counterstained with hematoxylin (Solarbio, G1080). Finally, slides were cover slipped and sealed with neutral resin. All clinical slides and tissue microarray slides (Tissue Array, HCol-Ade060CS1-01, HDgS-C140PT-01) were automatically scanned using a Leica Aperio Versa at 40x magnification. Scoring of PALMD staining was performed by an experienced pathologist with gastrointestinal tract expertise, using fraction/percentage of area (0.00 – 1.00) × intensity: 0 (Negative), 1+ (weakly positive), 2+ (intermediately positive), 3+ (strongly positive) in different cell types of colorectal tumor and adjacent normal tissues. For murine-derived samples, FFPE sections from 8 week old wild-type C57/B6 mice or excised SW480 oeCtrl and oePALMD tumors were processed and stained with primary antibody against PALMD (Proteintech, 16531-1-AP), R-spondin2 (Bioss, bs-18876R), LGR5 (Bioss, bs-20747R), or Lysozme (Abcam, ab108508) antibody as appropriate for immunohistochemistry. For normal mouse intestine and colon immunofluorescence staining, PALMD (Immunoway, YT7484) and Lysozyme (Abcam, ab108508) primary antibodies were used in combination with a 594/488 fluorescent dual-staining kit (Immunoway, RS0036). Immunofluorescence images were taken on a Stellaris 8 confocal microscope (Leica).

### Pharmacological Analysis Using the Cancer Therapeutics Response Portal (CTRP)

Proteomics and drug sensitivity data for colorectal adenocarcinoma cell lines were downloaded directly from CTRP (https://depmap.org/portal/interactive/)^42–44^. Specifically, the proteomic expression of PALMD was compared with the drug sensitivity data for emodin (PRISM Repurposing Primary Screen 19Q4) by Pearson’s correlation. Prior to correlation analysis the normality of the data was confirmed by the Kolmogorov-Smirnov test, and a single outlier cell line (Colo678) was detected and removed according to Dixon’s Q-test.

### TCGA RNA-Seq Analysis

Genomic Data Commons RNA-Sequencing data and associated clinical data for The Cancer Genome Atlas’ Colon Adenocarcinoma dataset (COAD) was downloaded from the UCSC cancer genome browser (xenabrowser.net) on December 30^th^, 2019. In total, RNA-Sequencing data was available for 453 primary tumor samples and 41 adjacent normal tissue samples. The intestinal stem cell (ISC) signature generated by the Batlle group was obtained directly from Table S1 of their publication^33^. Next, the edgeR package was used to identify genes strongly associated with the ISC signature, while controlling for the influence of disease stage. Candidate genes were manually checked against IHC staining in the Human Protein Atlas database for ISC-like localization to the crypt base of normal human intestinal tissues. Candidates with ISC-like localization were then checked for apparent overexpression in colorectal cancer compared to normal colon tissue. Candidates with apparent overexpression were checked for their association with patient outcomes in terms of progression-free survival (PFS) and cancer-specific survival (CSS) in the COAD dataset. Finally, TCGA’s colorectal adenocarcinoma (COADREAD) RNA-Seq dataset (Illumina HiSeqV2) was downloaded from the UCSC Cancer Genome Browser on May 22, 2020 and cross referenced with clinical chemotherapy data available from the BROAD GDAC FIREHOSE. Patients receiving first-line chemotherapy were manually determined from the dataset. Survival analyses were then performed for disease-free survival (DFS), PFS, and CSS based on these classifications. Samples were clustered into CRC consensus molecular subtypes (CMS) using the *CMSclassifier* R package^5^.

### Single-cell RNA-Seq Analysis

For t-SNE projections and visualization of PALMD and co-expressed markers, 10x cloupe files were obtained from the Human Colon Cancer Atlas (c295) dataset available at Broad’s Single Cell Portal (https://singlecell.broadinstitute.org/single_cell/study/SCP1162/human-colon-cancer-atlas-c295). The cloupe files were visualized using the Loupe Browser v6 software (10x Genomics) and clustered based on cell type and tissue type. For further analysis, the scRNA-Seq count data in H5 format was downloaded from the NCBI Gene Expression Omnibus (NCBI GEO), accession number GSE178341^55^. The h5 data was read into R v4.0.5 and converted into a CDS object for use in Monocle3^56–58^ analysis. The CDS was subsetted to include colorectal adenocarcinoma epithelial cells only and preprocessed using principal component analysis (PCA) with 50 dimensions using the *preprocess_cds* function. Following preprocessing, cells were aligned by batch ID (*align_cds*) and dimensionally reduced (*reduce_dimension* function) using the UMAP method^59^. UMAP projections were then plotted using the *plot_cells* function and colored by relevant characteristics (*e.g.* cell type or gene expression level). Pseudotime trajectory data was prepared using the *cluster_cells*, *learn_graph*, and *order_cells* functions. The primary lineage of interest for the target gene (CSC-CRC-tuft cell lineage) was selected for pseudotime analysis using the *choose_graph_segments* function. The relationship between genes of interest and pseudotime were plotted using the functions: *plot_cells* (UMAP plots) and *plot_genes_in_pseudotime* (expression vs. pseudotime plots).

### RNA-Sequencing of SW480 oeCtrl and oePALMD stable cell lines

SW480 oeCtrl and oePALMD stable cell lines were cultured in 100 mm^3^ cell culture dishes to 90% confluence, then washed gently with PBS 3 times and lysed into Trizol (Invitrogen, 15596018). Samples were shipped on dry ice to Allwegene Technology Inc (Beijing, CN) for RNA extraction and RNA-Seq. Upon arrival, RNA was extracted using standard Trizol isolation techniques and treated with DNAse I (Takara). RNA quality was assessed by gel electrophoresis and an Agilent 2100 Bioanalyzer (Agilent Technologies). The quantity and integrity of the RNA was determined using a NanoDrop spectrophotometer (Thermo). For library preparation and generation, 1.5 µg RNA from each sample was used as input, and then processed using an NEBNext® Ultra™ RNA Library Prep Kit for Illumina (NEB) according to the manufacturer instructions. Next, an AMPure XP System was used to filter library fragments of appropriate size (200-200bp). Following PCR, products were again assessed for quality using an Agilent 2100 Bioanalyzer. Finally, the library fragments were sequenced using the Illumina NovaSeq 6000 platform, and paired-end 150 bp reads were generated. For quality control, low quality reads, reads containing adapter, and reads containing poly-N reads were removed from the raw data. Clean reads were then mapped to the reference human genome using STAR. HTSeq v0.5.4 p3 was used to count the reads mapped to each gene. Differential expression analysis was performed using GSEA. V4.2.3 and a heatmap was generated using the *heatplot* function in the made4 package in R v4.0.5.

### Xenograft experiments

SW480 oeCtrl or oePALMD stable cells were injected subcutaneously into the flanks of 6-week old athymic nude mice at 1 × 10^6^ cells per mouse (n=10 and n=11 respectively). Mice were housed in a specific pathogen-free conditions with free access to food and water. Flanks were assessed every 2 days for tumor growth and mouse weights were monitored. Upon detection of tumors, measurements were taken with a caliper every 2 days. One oeCtrl mouse experienced weight loss, failed to form a tumor, died for unknown reasons, and therefore was excluded from analysis. Tumor volumes were calculated based on the formula V=(L x W^2^)/2. On the final day of the experiment, mice were euthanized; tumors were excised, measured, weighed, and separated for Western blotting, ELISA, and immunohistochemistry.

### Statistical Analysis

Statistical analyses were performed using GraphPad Prism 8 (GraphPad Software, San Diego, CA, USA). Significant differences between the mean values of each group were detected using the Student’s t-test. For non-parametric comparisons, the Mann-Whitney U test was performed. P values <0.05 were considered statistically significant. For correlation analysis the Pearson method was used and Kendall’s correlation rank tau was determined in R using the *cor.test* function. For survival analyses, the Cox proportional hazards model and the log-rank test were used where appropriate. Survival plots were generated using the survminer R package and cutpoints were determined using the *surv_cutpoint* function based on maximally-ranked statistics.

## Data Availability

RNA-Sequencing raw data for SW480 oeCtrl and oePALMD stable cell lines is available through NCBI Sequencing Read Archive (PRJNA1110604). Human Colon Cancer Atlas scRNA-Seq data is available from NCBI GEO accession no. GSE178341. TCGA COAD and COADREAD data are available at xenabrowser.net. Cell sensitivity data are available from the Cancer Therapeutics Response Portal at https://depmap.org/portal/interactive. All other data is available upon reasonable request from the corresponding authors.

## Author Contributions

YN Y, JS, and NW designed the initial study; YN Y, JS, YP Y, and NW performed patient-derived organoid studies; TY and ZY provided surgical/biopsy samples, blood serum, and obtained clinical information of CRC patients; SL, RY, JY, and LC performed immunohistochemistry and immunofluorescence studies and pathological assessment; NW and JY performed bioinformatic and clinical analyses; YN Y, YP Y, YL, LF, LD, HC, QL, ZZ, and NW performed in vitro functional and molecular experiments; SL, HF, and NW performed in vivo experiments and analysis; PC, DQ, and JD provided experimentation advice; JD, ZC, JP, and NW provided funding and materials; YN Y, JS, ZC, JP, and NW wrote the initial manuscript; PC, DQ, JD, ZC, JP, and NW revised the manuscript; ZC, JP, and NW supervised the project; All authors reviewed and approved the manuscript.

