## Supplementary material for "Palmdelphin facilitates R-spondin2 secretion to activate Wnt signaling and promote colorectal cancer stemness and tumorigenesis": Figures S1-9

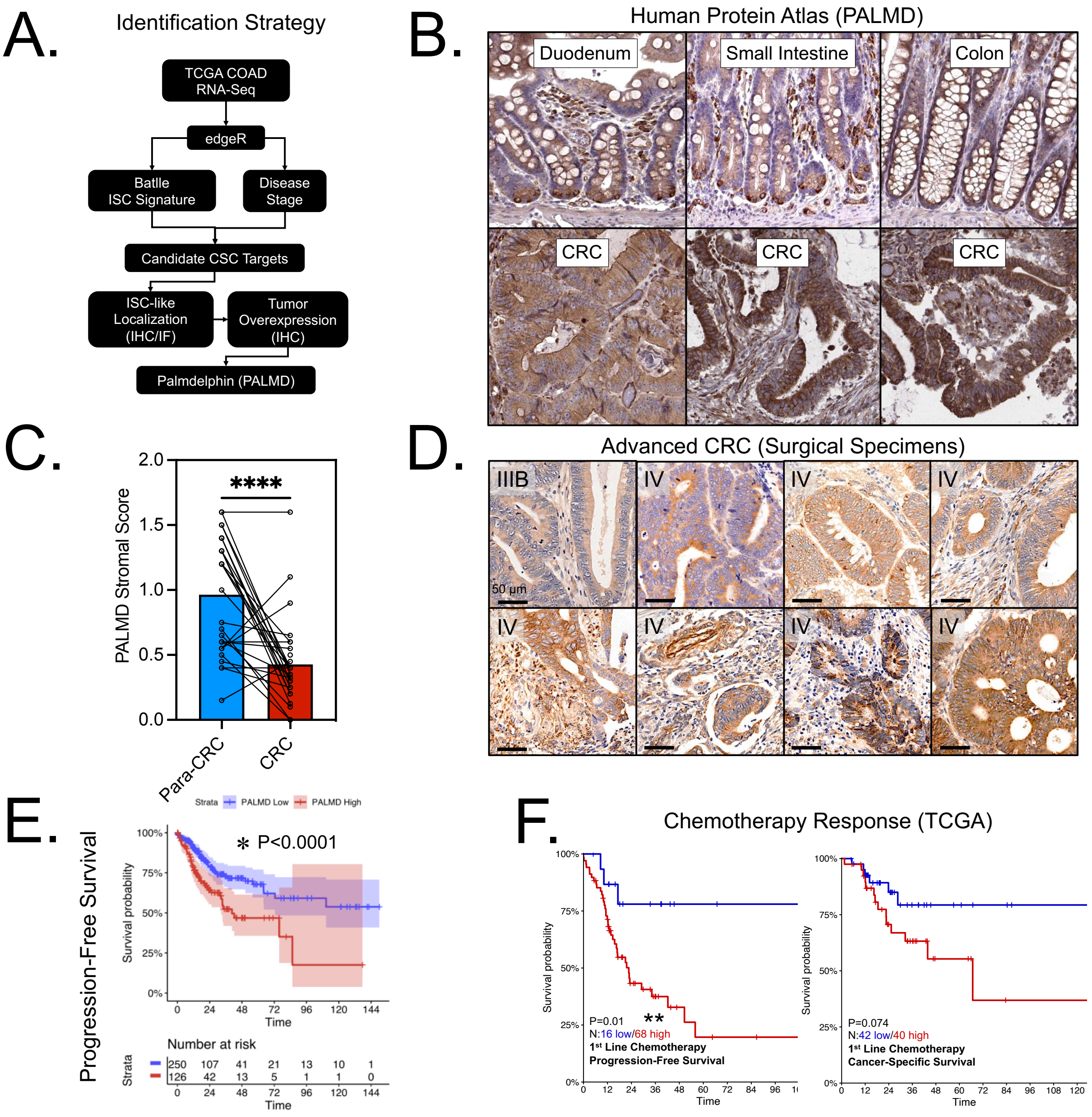


**Figure S1. Additional pathological information for PALMD in CRC. A.** Flowchart depicting candidate novel CSC target identification strategy. Analysis was performed using the Batlle stem cell signature and disease stage in edgeR to interrogate the TCGA's colon adenocarcinoma dataset, which was followed by confirmation of ISC-like localization and tumor overexpression by immunohistochemistry. **B.** Immunohistochemical staining from the human protein atlas demonstrating PALMD strongly positive cells interspersed at the base of intestinal epithelial crypt.

**C.** Comparison of pathologist stromal scoring (intensity x area) for Palmdelphin in CRC compared to para-CRC showing a significant decrease in CRC (paired T-test P<0.0001). **D.** Immunohistochemical staining of PALMD in Stage IIIB-IV CRC surgical specimens demonstrating frequent epithelial expression ranging from moderate and diffuse to strong and specific, with frequent cytoplasmic and occasional membrane-like localization. **E.** High expression levels of PALMD are associated with earlier progression onset in the TCGA colorectal cancer dataset (log-rank test P<0.0001). **F.** Survival analysis for CRC patients receiving adjuvant or first-line chemotherapy demonstrating a significant increase in progression in first-line chemotherapy patients expressing high levels of PALMD.

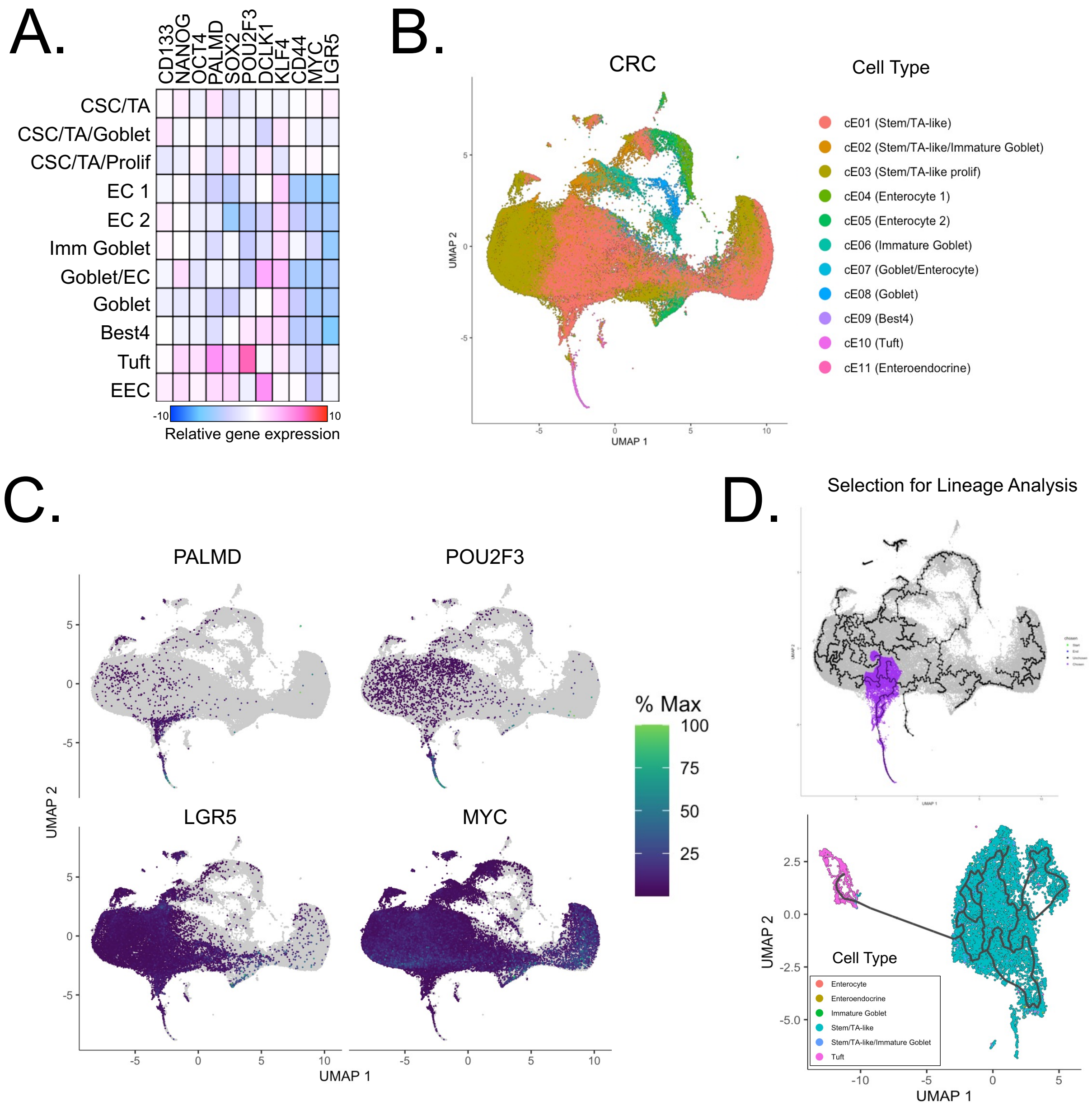


**Figure S2. scRNA-Seq UMAP projection of PALMD and select CSC and tuft cell markers. A.** Coexpression plot for PALMD and selected stem cell (LGR5, CD44, MYC/c-MYC, PROM1/CD133, POU2F1/OCT4, NANOG, KLF4, SOX2) and tuft cell (DCLK1, POU2F3) genes revealing coexpression for: PALMD, DCLK1, and POU2F3 in normal tuft cells; PALMD, POU2F3, SOX2, NANOG, OCT4, and KLF4 in tumor tuft cells; and PALMD, LGR5, and c-MYC in tumor stem/TA-like cells (CSCs) in the Human Colon Cancer Atlas (HCCA) single cell RNA-Sequencing dataset (GSE178341). **B.** UMAP projection for single-cell RNA sequencing data obtained from the human colon cancer atlas and filtered to only include colon adenocarcinoma, showing lineage relationships between epithelial cell types. **C.** UMAP expression plot for PALMD, POU2F3, LGR5, and MYC showing prominent PALMD expression in an apparent CSC-to-tumor tuft cell lineage. D. Trajectory map of UMAP projection of human colon cancer atlas colon adenocarcinomas, showing the lineage/cells selected for further analysis.

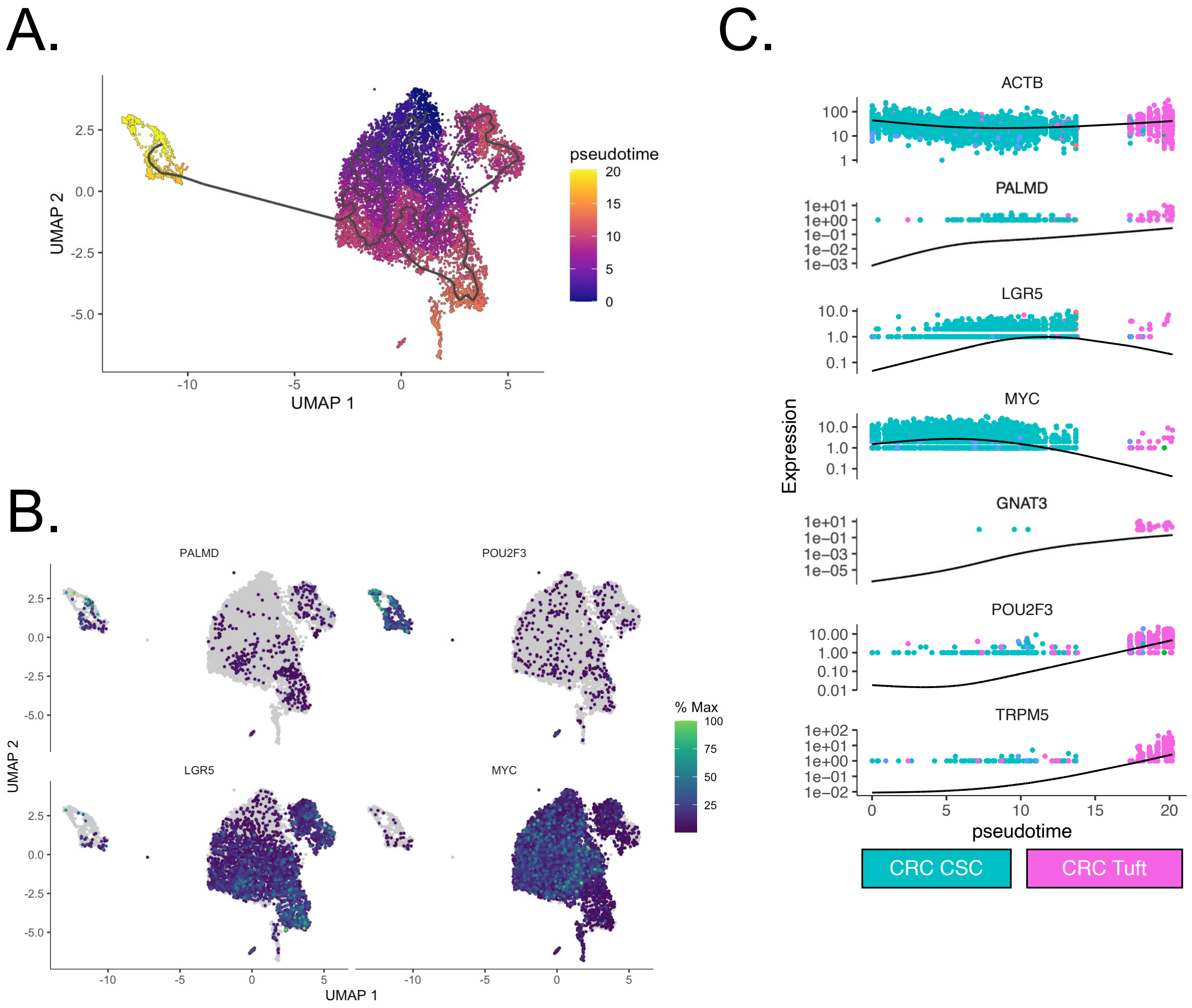


**Figure S3. Palmdelphin expression is consistent with a CSC/tuft cell lineage in single-cell RNA-Seq pseudotime trajectory analysis. A.** Pseudotime UMAP trajectory plot of the selected CSC-to-tumor tuft cell lineage showing hypothetical tuft cell differentiation from CSCs. **B.** UMAP expression plots for PALMD, POU2F3, LGR5, and MYC demonstrating prominent PALMD expression in the CSC-to-tumor tuft cell lineage. **C.** Gene expression plots for β-actin (ACTB - control), PALMD, CSC markers (LGR5, MYC) and tuft cell markers (GNAT3, POU2F3, TRPM5) demonstrating both CSC and tumor tuft-like expression dynamics for PALMD across pseudotime.

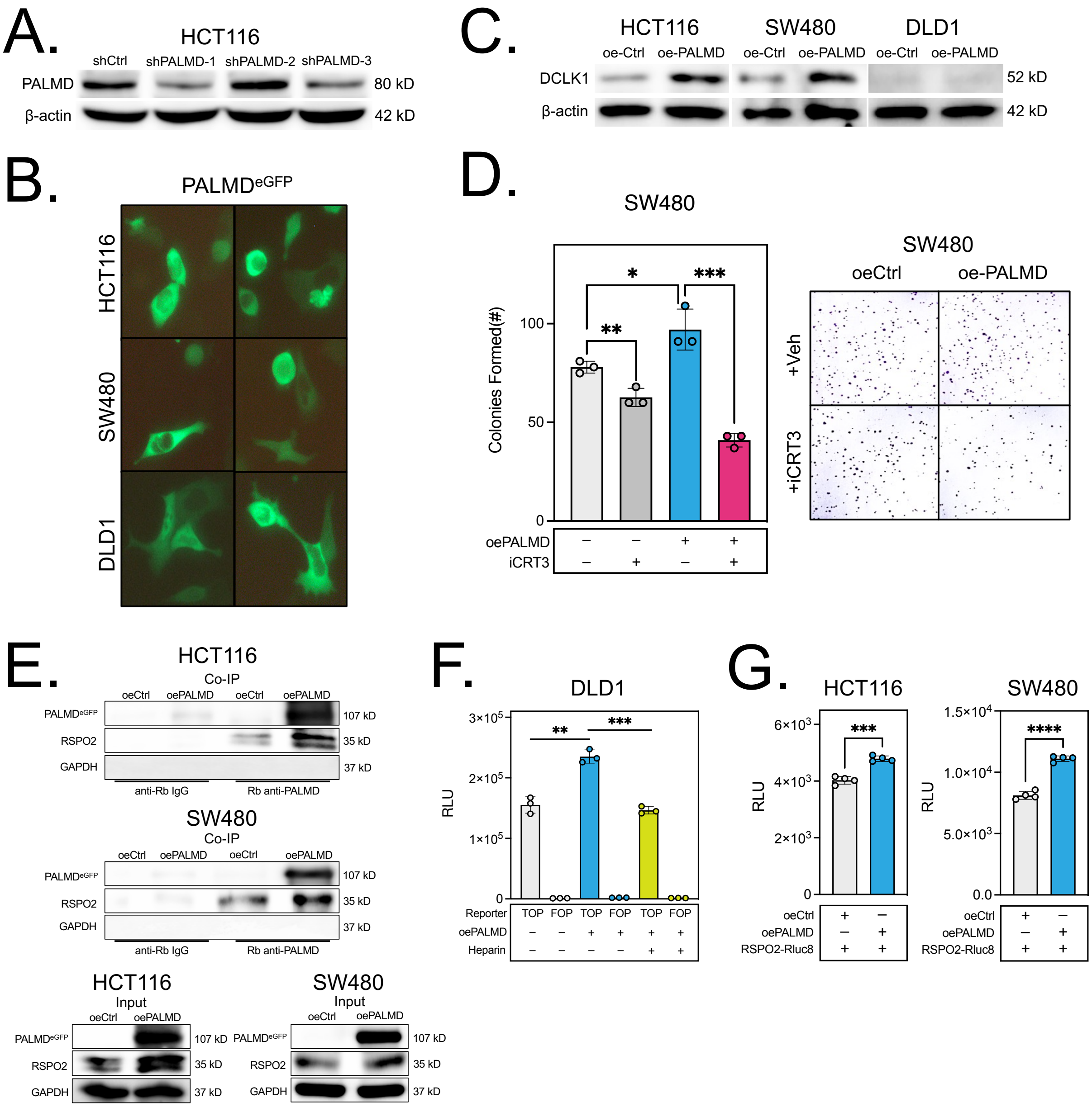


**Figure S4. Additional information for PALMD expression and its effect on Wnt signaling pathway. A.** Western blotting results for 3 PALMD shRNAs in HCT116 cells transfected for 48 h demonstrating equal effectiveness for sh-1 and sh-3 (selected). **B.** Fluorescence microscopy of PALMD$^{eGFP}$ fusion protein expression in HCT116, SW480, and DLD1 cells. **C.** DCLK1 expression is increased after overexpression of PALMD in HCT116 and SW480 cells. **D.** Quantification of colony numbers in iCRT3-treated SW480 cells revealing that PALMD-promoted CRC colony formation is attenuated by Wnt pathway inhibition. **E.** Co-immunoprecipitation results from

HCT116 and SW480 cell lines overexpressing PALMD reveal an increase in the PALMD/R-spondin2 interaction in oePALMD compared to oeCtrl. **F.** Heparin treatment significantly attenuates TOPFlash reporter activation in DLD1 cells overexpressing PALMD. **G.** Quantification of *Renilla* luciferase reporter expression in culture medium from HCT116 and SW480 cells co-transfected with oeCtrl or oePALMD and RSPO2-RLuc8 plasmid, revealing a statistically significant increase in the medium of oePALMD/RSPO2-Luc8 co-tranfected cells compared to controls. *P<0.05, **P<0.01, ***P<0.001.

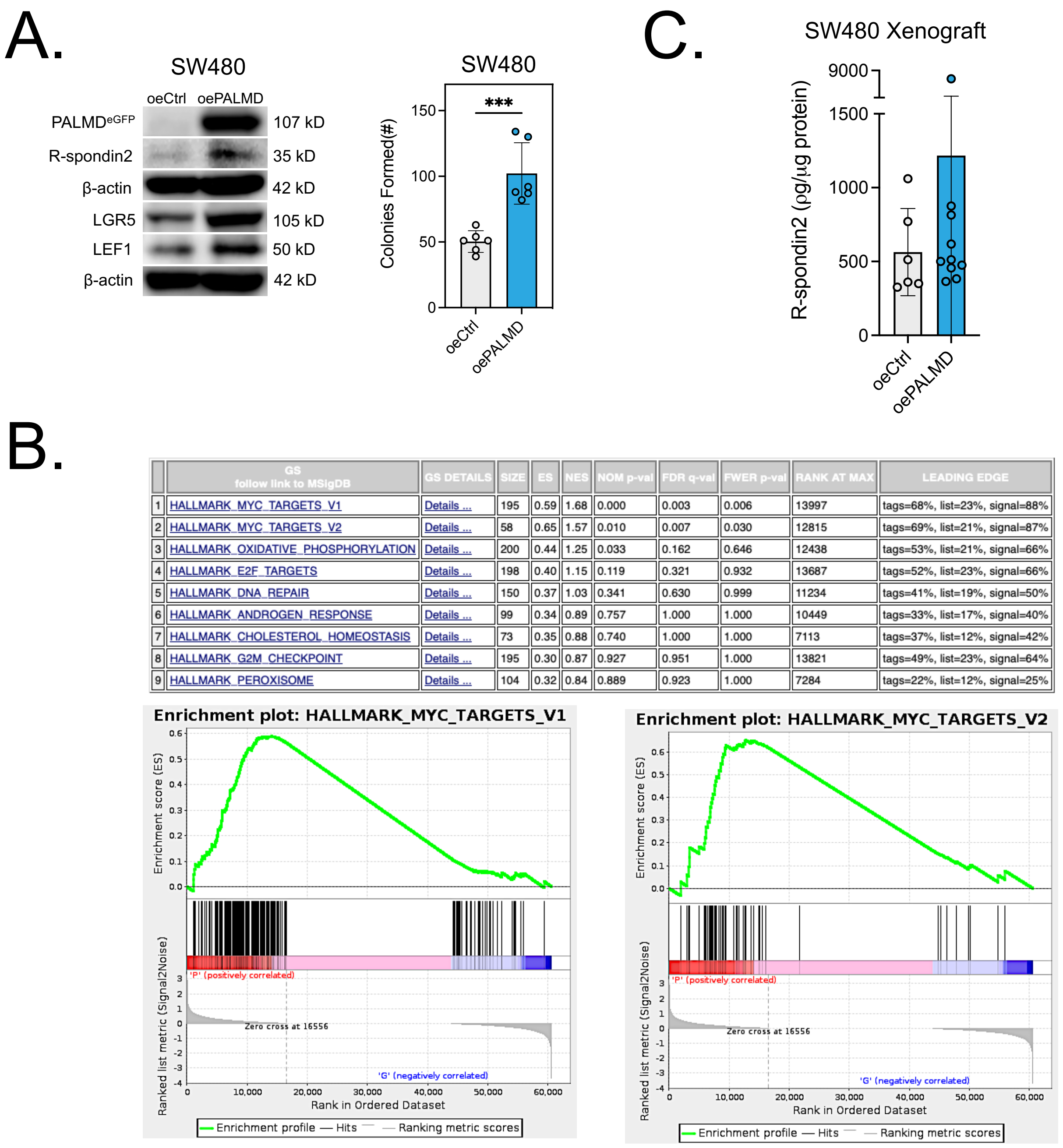


| | GS follow link to MSigDB | GS DETAILS | SIZE | ES | NES | NOM p-val | FDR q-val | FWER p-val | RANK AT MAX | LEADING EDGE |
|---|---|---|---|---|---|---|---|---|---|---|
| 1 | HALLMARK_MYC_TARGETS_V1 | Details ... | 195 | 0.59 | 1.68 | 0.000 | 0.003 | 0.006 | 13997 | tags=68%, list=23%, signal=88% |
| 2 | HALLMARK_MYC_TARGETS_V2 | Details ... | 58 | 0.65 | 1.57 | 0.010 | 0.007 | 0.030 | 12815 | tags=69%, list=21%, signal=87% |
| 3 | HALLMARK_OXIDATIVE_PHOSPHORYLATION | Details ... | 200 | 0.44 | 1.25 | 0.033 | 0.162 | 0.646 | 12438 | tags=53%, list=21%, signal=66% |
| 4 | HALLMARK_E2F_TARGETS | Details ... | 198 | 0.40 | 1.15 | 0.119 | 0.321 | 0.932 | 13687 | tags=52%, list=23%, signal=66% |
| 5 | HALLMARK_DNA_REPAIR | Details ... | 150 | 0.37 | 1.03 | 0.341 | 0.630 | 0.999 | 11234 | tags=41%, list=19%, signal=50% |
| 6 | HALLMARK_ANDROGEN_RESPONSE | Details ... | 99 | 0.34 | 0.89 | 0.757 | 1.000 | 1.000 | 10449 | tags=33%, list=17%, signal=40% |
| 7 | HALLMARK_CHOLESTEROL_HOMEOSTASIS | Details ... | 73 | 0.35 | 0.88 | 0.740 | 1.000 | 1.000 | 7113 | tags=37%, list=12%, signal=42% |
| 8 | HALLMARK_G2M_CHECKPOINT | Details ... | 195 | 0.30 | 0.87 | 0.927 | 0.951 | 1.000 | 13821 | tags=49%, list=23%, signal=64% |
| 9 | HALLMARK_PEROXISOME | Details ... | 104 | 0.32 | 0.84 | 0.889 | 0.923 | 1.000 | 7284 | tags=22%, list=12%, signal=25% |

**Figure S5. Additional data for SW480 stable cell lines. A.** Immunoblotting and colony formation assay results confirming the molecular (R-spondin2/Wnt protein increased expression) and functional characteristics (increased colony formation) of the stable SW480 oePALMD cell line. **B.** Gene enrichment analysis results (Hallmark-50) for the stable SW480 oePALMD cell line identifying MYC targets as the top 2 enriched pathways. **C.** R-spondin2 concentration in SW480 xenograft tumor lysates as determined by ELISA, demonstrating an approximately two-fold average increase (ns) in oePALMD tumors compared to control. ***P<0.001.

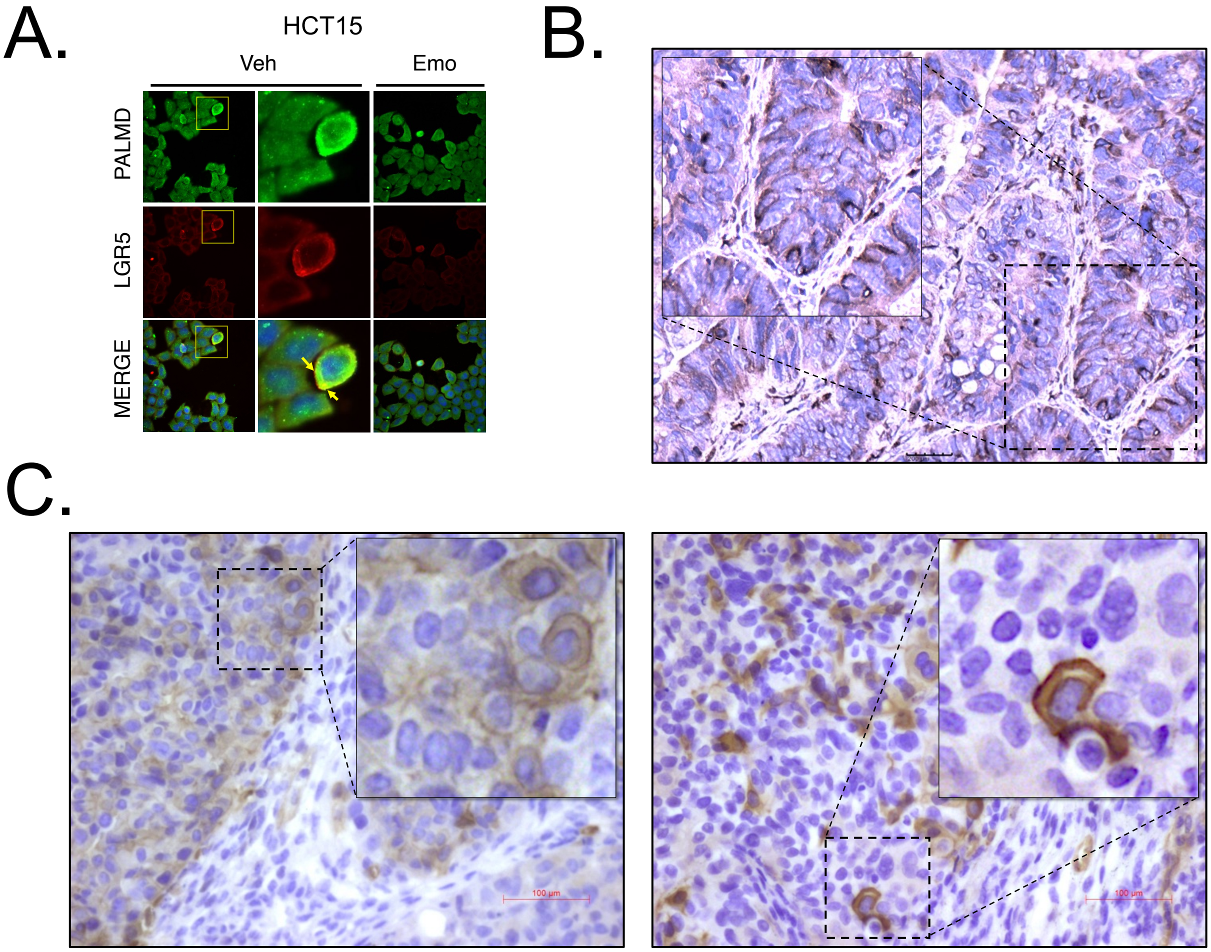


**Figure S6. Additional evidence for membrane localization of PALMD. A.** Immunofluorescence staining for PALMD and LGR5 in HCT15 cells after 48 h of treatment with vehicle (Veh) or emodin (Emo; 10 μM) demonstrating PALMD/LGR5 membrane co-localization and significantly decreased expression after treatment. **B.** Immunohistochemical staining from CRC patient tumor tissue demonstrating potential cytoplasmic, perinuclear, and membrane localization of PALMD. **C.** Immunohistochemical staining from oePALMD xenograft tissue demonstrating apparent membrane localization of PALMD.

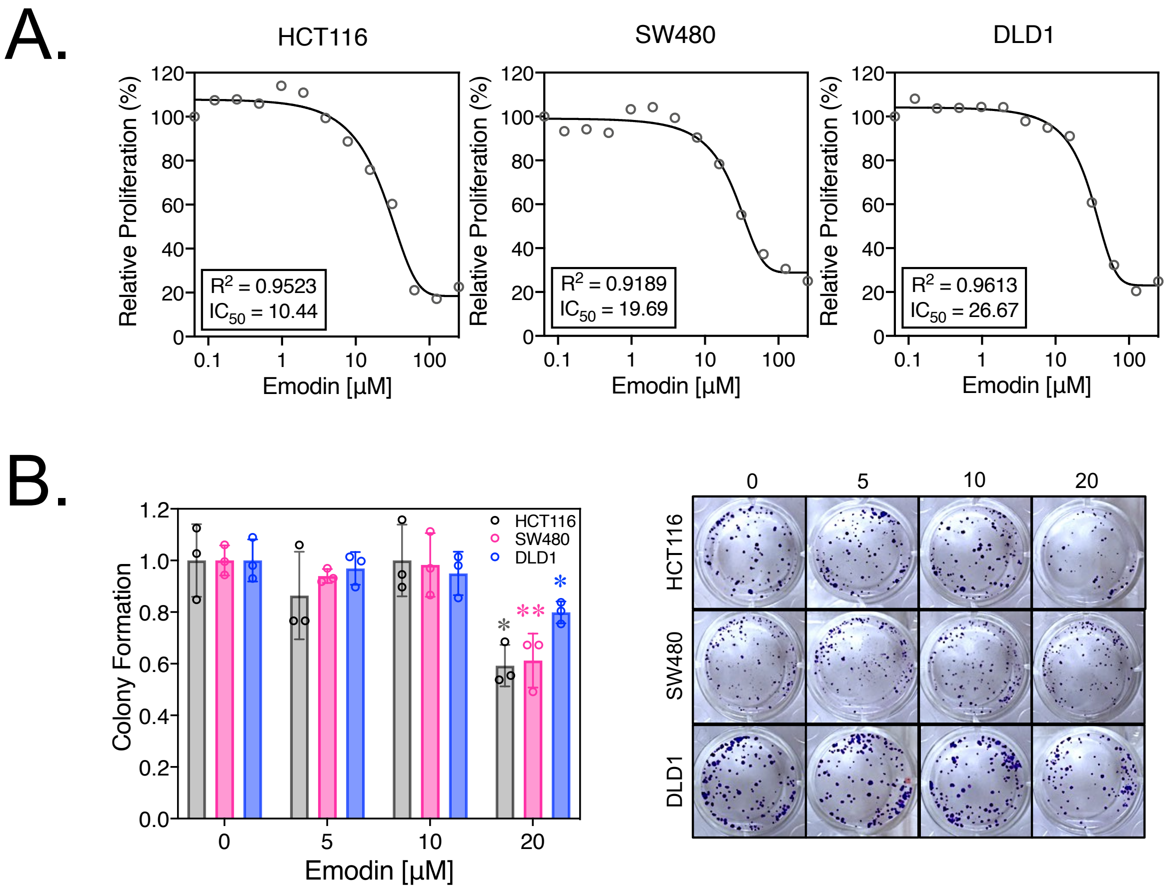


**Figure S7. Emodin has weak anti-proliferative/cytotoxic activity in CRC cell lines. A.** CCK8 assay results demonstrating IC50 values ranging from 10.44 – 26.67 in HCT116, SW480, and DLD1 cells after 48 h of emodin treatment. **B.** Quantification of colony formation results demonstrating no significance for emodin at concentrations less than 20 µM. **C.** Representative images of the colony formation assay quantified in B. *P<0.05, **P<0.01.

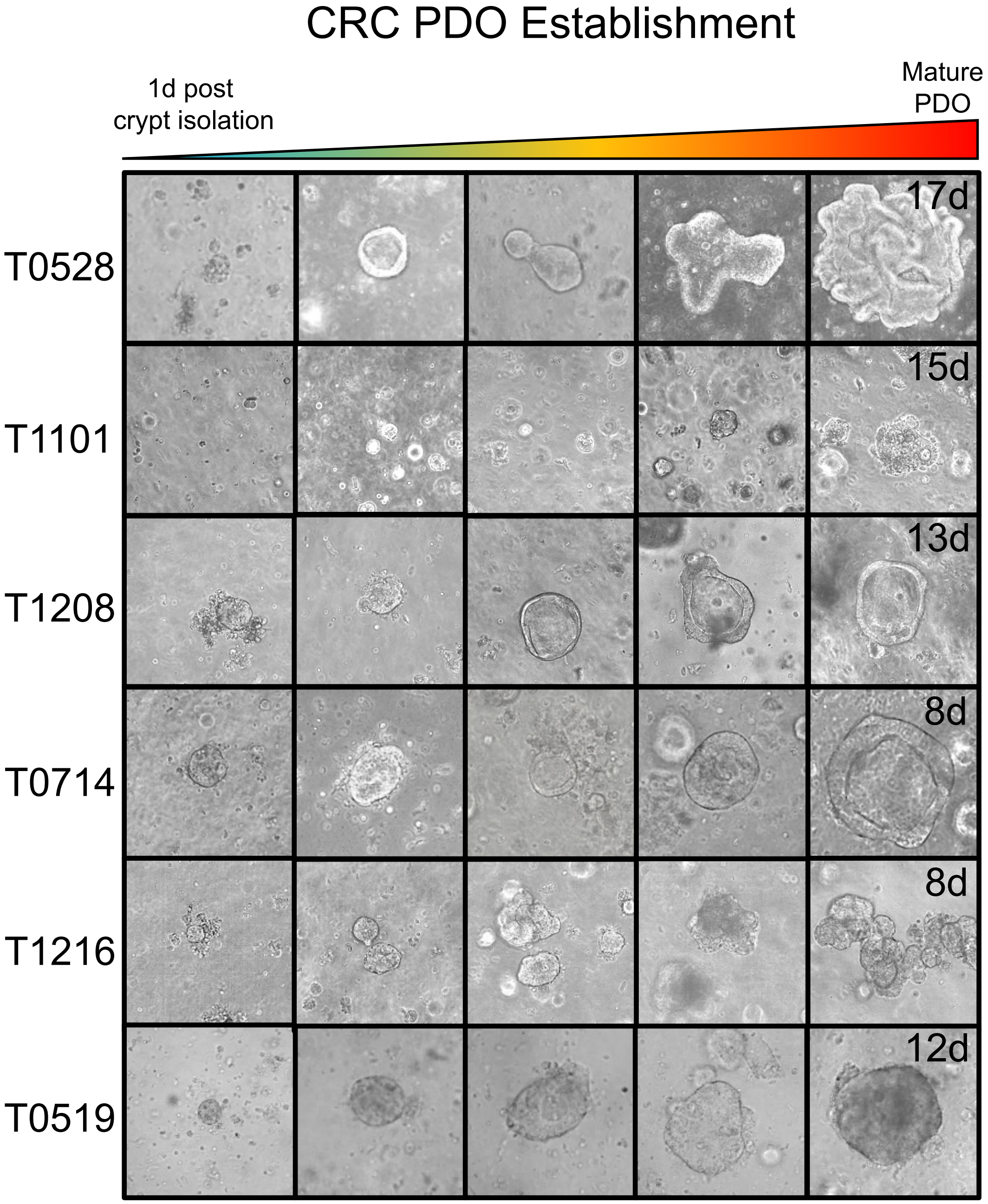


**Figure S8. Representative figure showing CRC PDO establishment from isolated epithelial crypts to mature organoids.**

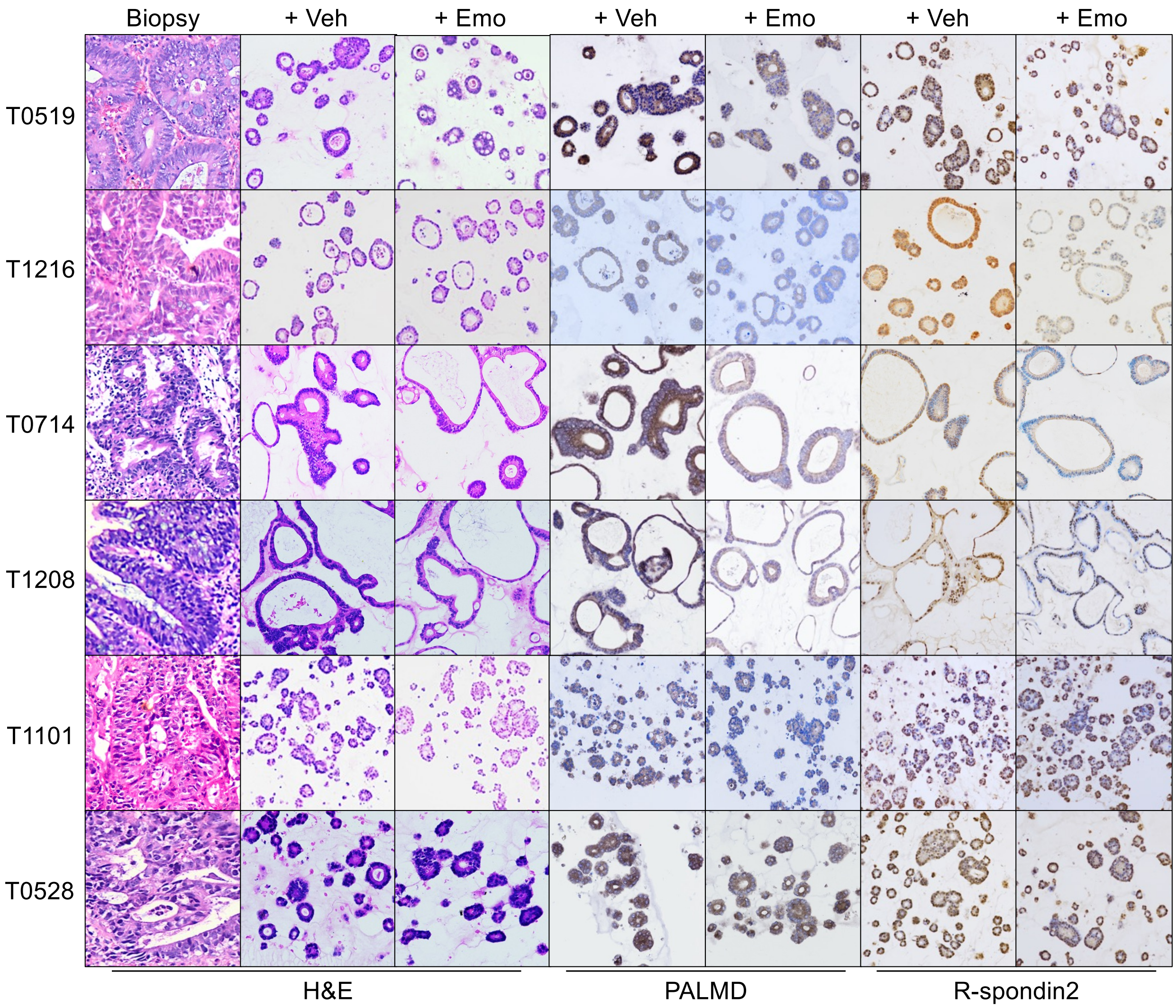


**Figure S9. Hematoxylin/Eosin and immunohistochemistry staining of patient biopsies compared to patient-derived organoids treated with vehicle or 10 µM of emodin.**
