## Supplemental Table I for "Palmdelphin facilitates R-spondin2 secretion to activate Wnt signaling and promote colorectal cancer stemness and tumorigenesis"

### Supplemental Table I. Patient characteristics for serum R-Spondin2 level versus tumor IHC comparisons

| ID | Sex | Age (Yrs) | AFP | CA-125 | CEA | CA19-9 | Anatomic Location | Pathology | T | N | M | Perineural Invasion | Vascular Embolism | Serum R-spondin2 (pg/ml) | PALMD IHC Score | R-spondin2 IHC Score | Nuclear Beta-Catenin IHC Score |
|---|---|---|---|---|---|---|---|---|---|---|---|---|---|---|---|---|---|
| P1 | Female | 70 | 1.13 | 11.6 | 11.95 | 4.55 | Ascending Colon | Moderately Differentiated Tubular Adenocarcinoma | 3 | 1 | 0 | Yes | Yes | 163.782 | 4 | 2 | 1 |
| P2 | Female | 72 | 2.43 | Not tested | 2.66 | 2.43 | Rectum | Signet Ring Cell Carcinoma with Mucinous Adenocarcinoma Components | 3 | 0 | 0 | Yes | No | 154.556 | 2 | 3 | 2 |
| P3 | Female | 77 | 1.15 | Not tested | 2.68 | 3.55 | Sigmoid Colon | Moderately Differentiated Tubular Adenocarcinoma | 3 | 0 | 0 | Yes | Yes | 222.845 | 9 | 9 | 6 |
| P4 | Female | 75 | 1.85 | 36.7 | 2.58 | 40.9 | Rectum | Moderately Differentiated Tubular Adenocarcinoma | 3 | 1 | 0 | Yes | Yes | 72.166 | 3 | 3 | 1 |
| P5 | Female | 59 | 3.71 | 10.1 | 0.8 | 2 | Rectum | Moderately Differentiated Tubular Adenocarcinoma | 2 | 1 | 0 | No | Yes | 60.401 | 2 | 3 | 0 |
| P6 | Male | 80 | 1.91 | Not tested | 2.09 | 2.58 | Sigmoid Colon | Moderately Differentiated Tubular Adenocarcinoma | 3 | 0 | 0 | No | No | 108.181 | 2 | 1 | 2 |
| P7 | Male | 67 | Not tested | Not tested | Not tested | Not tested | Sigmoid Colon | Moderately Differentiated Tubular Adenocarcinoma | 2 | 0 | 0 | No | No | 261.547 | 6 | 3 | 2 |
| P8 | Male | 44 | 2.76 | Not tested | 35.78 | 1434.23 | Rectum | Moderately Differentiated Tubular Adenocarcinoma | 3 | 1 | 0 | Yes | Yes | 151.115 | 2 | 1 | 1 |
| P9 | Male | 64 | 3.38 | Not tested | 5.19 | 19.93 | Rectum | Moderately to Poorly Differentiated Adenocarcinoma | 3 | 1 | 0 | Yes | Yes | 156.512 | 4 | 3 | 2 |
| P10 | Female | 54 | 2.06 | Not tested | Not tested | 4.15 | Ileocecal Junction | Moderately Differentiated Tubular Adenocarcinoma | 3 | 1 | 0 | Yes | Yes | 97.169 | 3 | 2 | 1 |
| P11 | Male | 61 | 2.65 | 45.5 | 50.24 | 56.18 | Sigmoid Colon | Moderately Differentiated Tubular Adenocarcinoma | 3 | 1 | 0 | Yes | Yes | 102.457 | 3 | 4 | 4 |
| P12 | Male | 62 | 3.27 | Not tested | 59.94 | 95.65 | Rectum | Moderately Differentiated Tubular Adenocarcinoma | 3 | 0 | 0 | Yes | Yes | 107.615 | 4 | 2.5 | 1 |
